# Intron-targeted mutagenesis reveals roles for *Dscam1* RNA pairing-mediated splicing bias in neuronal wiring

**DOI:** 10.1101/622217

**Authors:** Weiling Hong, Haiyang Dong, Jian Zhang, Fengyan Zhou, Yandan Wu, Yang Shi, Shuo Chen, Bingbing Xu, Wendong You, Feng Shi, Xiaofeng Yang, Zhefeng Gong, Jianhua Huang, Yongfeng Jin

## Abstract

*Drosophila melanogaster* Down syndrome cell adhesion molecule (*Dscam1*) can potentially generate 38,016 different isoforms through stochastic, yet highly biased, alternative splicing. Genetic studies demonstrated that stochastic expression of multiple Dscam1 isoforms provides each neuron with a unique identity for self/non-self-discrimination. However, due to technical obstacles, the functional significance of the highly specific bias in isoform expression remains entirely unknown. Here, we provide conclusive evidence that *Dscam1* splicing bias is required for precise mushroom body (MB) axonal wiring in flies in a variable exon-specific manner. We showed that targeted deletion of the intronic docking site perturbed base pairing-mediated regulation of inclusion of variable exons. Unexpectedly, we generated mutant flies with normal overall Dscam1 protein levels and an identical number but global changes in exon 4 and exon 9 isoform bias (DscamΔ4D^−/−^ and DscamΔ9D^−/−^), respectively. DscamΔ9D^−/−^ mutant exhibited remarkable mushroom body defects, which were correlated with the extent of the disrupted isoform bias. By contrast, the DscamΔ4D^−/−^ animals exhibited a much less severe defective phenotype than DscamΔ9D^−/−^ animals, suggestive of a variable domain-specific requirement for isoform bias. Importantly, mosaic analysis revealed that changes in isoform bias caused axonal defects but did not influence the self-avoidance of axonal branches. We concluded that, in contrast to the Dscam1 isoform number that provides the molecular basis for neurite self-avoidance, isoform bias may play a non-repulsive role in mushroom body axonal wiring.

## Introduction

Neural circuits are precisely assembled through interactions among neurites of enormous numbers of neurons^1^. This assembly relies upon diverse cell surface receptors mediating repulsive and adhesive interactions between neurites, including the *Drosophila* Down syndrome cell adhesion molecule (*Dscam1*) and the mammalian clustered protocadherins (*PCDH*s)^2^. *Drosophila Dscam1* can potentially generate 38,016 different isoforms through alternative splicing, and has been suggested as an important regulator of neural circuits^3^. The Dscam1 isoforms have been shown to exhibit isoform-specific homophilic binding almost exclusively, with little or no heterophilic binding^4, 5^. Furthermore, each neuron acquires a unique cell surface identity through expression of a unique combination of tens of isoforms in a largely stochastic manner^6, 7, 8^. The stochasticity in the generation of Dscam1 isoform repertoires enables each neuron to discriminate itself from all of the other cells. Genetic studies revealed the importance of Dscam1 for self-avoidance and normal patterning of axons and dendrites^7, 9, 10, 11, 12, 13, 14, 15, 16, 17^. Deletion mutants combined with mathematical modeling suggested that thousands of Dscam1 isoforms are required for discrimination of self versus non-self dendrites^18^. Taken together, these results indicate that Dscam1 diversity is required for self-recognition and neuronal wiring.

However, most previous functional experiments focused mainly on the role of *Dscam1* itself and its number of isoforms. In contrast, few studies have examined the effects of isoform bias on neuronal wiring. This is more relevant to our understanding of the function of Dscam1 isoform diversity because the isoforms from exon 4, 6, and 9 clusters are highly regulated in a tissue- or cell type-specific manner^6, 7, 8, 19, 20, 21^. This observation suggests that Dscam1 isoform profiles arise from a series of stochastic but biased splicing events in individual cells. Indeed, analyses of alternative exon 4 expression using splicing reporters indicated that *Dscam1* splicing is probabilistic, which is compatible with a canonical role in neural circuit assembly through self-avoidance^8^. However, the role of biased splicing obviously cannot be explained by this self-avoidance model. In contrast, biased splicing seems not to be economical for neurite self-avoidance, because highly biased splicing heavily limits access to the full diversity from a pool of Dscam1 isoforms^18, 22^. Although recent studies suggested that biased isoform expression may be combinatorially regulated by the strength of base pairing between the docking site and selector sequence, the strength of the splice sites, and RNA binding proteins^23, 24, 25, 26, 27, 28^, the physiological significance of the highly specific bias in isoform expression remains entirely unclear.

If the specific isoform bias plays a crucial role in neuronal development, then disruption of that bias should cause a phenotypic alteration in neurons. However, isoform bias-disrupting experiments are hampered by significant technical obstacles, as excessive or reduced Dscam1 protein levels are likely to cause phenotypic defects^3, 7, 13, 17, 29, 30^. Previous functional experiments using knock-ins, genomic deletions, or isoform overexpression have either resulted in altered Dscam1 protein levels, a reduced number of isoforms, or both. Therefore, it remains unclear whether these phenotypic defects observed in *Dscam1* mutants are caused by changes in the number of isoforms, isoform bias, and/or protein levels. A definitive test to address the importance of the Dscam1 isoform bias would involve generation of mutant flies with dynamically changing isoform ratios, but with identical protein levels and the potential for isoform numbers similar to those observed in wild type controls.

Here, we provide conclusive evidence that *Dscam1* splicing bias is required for precise mushroom body (MB) axonal wiring in a variable exon-specific manner. We showed that deletion of the intronic docking site perturbed base pairing-mediated regulation of variable exon inclusion of *Dscam1*. Unexpectedly, we generated viable mutant flies with normal overall Dscam1 protein levels and an identical number of Dscam1 isoforms, albeit showing a change in in exon 4 and exon 9 isoform bias (DscamΔ4D^−/−^ and DscamΔ9D^−/−^). DscamΔ9D^−/−^ mutant exhibited remarkable mushroom body defects, which were correlated with the extent of the disrupted isoform bias. By contrast, the DscamΔ4D^−/−^ animals exhibited a much less severe defective phenotype than DscamΔ9D^−/−^ animals, suggestive of a variable domain-specific requirement for isoform bias. These studies indicated that not only a very large number, but also a bias of Dscam1 isoforms, are necessary to ensure normal neuronal wiring. Our findings suggest a distinct role for Dscam1 isoforms in neural circuitry beyond canonical self-avoidance.

## Results

### Deletion of the docking site of exon cluster 4 or 9 did not affect overall variable exon inclusion or Dscam1 protein level

The initial aim of the present study was to elucidate the functional roles of the docking site in regulating *Dscam1* alternative splicing in *Drosophila* using the CRISPR/Cas9 system, because the *Dscam1* exon 9 minigene did not work in transfection experiments using *Drosophila* S2 cells. To address this issue, we deleted the docking site of the exon 9 cluster by CRISPR/Cas9, and thus generated homozygous viable flies lacking the docking site of the exon 9 cluster (designated DscamΔ9D^−/−^) (Fig. 1a; Supplementary Fig. 1). DscamΔ9D^−/−^ adults did not show visible phenotypic defects in comparison with wild type controls. Surprisingly, reverse transcription PCR (RT-PCR) using primers aligning to exons 8 and 10 of *Dscam1* demonstrated that there was no exon 9 skipping in the head and other tissues of DscamΔ9D^−/−^ flies, similar to the wild type controls (Fig. 1b). Further quantitative real-time PCR revealed that the overall amount of *Dscam1* transcripts was indistinguishable between DscamΔ9D^−/−^ mutants and wild type controls (Fig. 1c). Moreover, the level of Dscam1 protein was indistinguishable between DscamΔ9D^−/−^ and wild type controls, as shown by Western blotting analysis. Taken together, these observations indicated that DscamΔ9D^−/−^ flies exhibited normal overall *Dscam1* expression levels similar to wild type controls.

**Figure 1.**
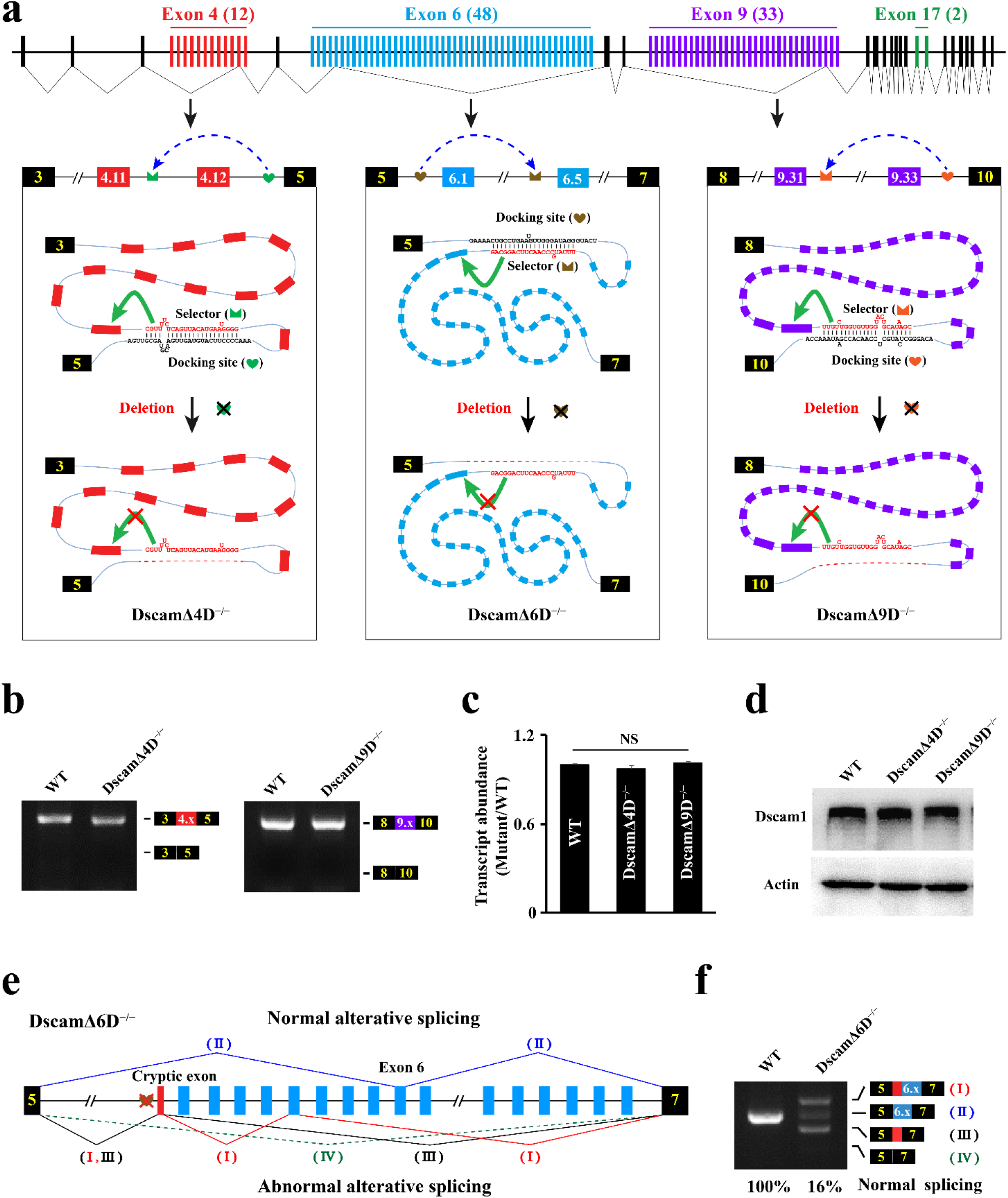
Generation and molecular characterization of DscamΔ4D^−/−^ and DscamΔ9D^−/−^ alleles. (**a**) Schematic diagrams of the docking site mutations in *D*. *melanogaster Dscam1*; docking sites (marked by hearts) were paired with the selector sequence (marked by crowns). The docking sites in the exon 4, 6, and 9 clusters were deleted; a green arrow indicates activation of the inclusion of the proximal exon and an arrow plus a black cross mark indicates the abolishment of the inclusion. (**b**) The exon 4 and exon 9 inclusions for each *Dscam1* mutant were indistinguishable from the wild type. (**c**) Deletion of the docking sites did not affect the levels of *Dscam1* transcripts. (**d)** Disruption of the docking sites did not affect the levels of Dscam1 protein. **(e)** Schematic diagrams of the mis-splicing caused by deletion of the docking site of exon cluster 6. An apparent cryptic exon (red box) is located immediately downstream of the docking site. Deletion of the docking site activated the cryptic exon splicing to variable exon 6 or constitutive exon 7, and exon 6 skipping. (**f)** Deletion of the docking site of exon cluster 6 markedly decreased normal exon 6 inclusion. The larger band than normal exon 6 inclusion corresponded to the abnormal product of cryptic exon splicing to variable exon 6 (I), while the smaller abnormal band corresponded to the product of cryptic exon splicing to constitutive exon 7 (III).

Using a similar strategy, we generated homozygous viable flies with deletion of the intronic docking site of exon 4 cluster (DscamΔ4D^−/−^, Fig. 1a; Supplementary Fig. 1). DscamΔ4D^−/−^ also showed no visible phenotypic defects. We did not detect any significant differences in exon 4 skipping between DscamΔ4D^−/−^ and wild type flies (Fig. 1b). Similar to DscamΔ9D^−/−^ mutants, overall *Dscam1* transcript and protein levels in DscamΔ4D^−/−^ flies were identical to those in wild type controls (Fig. 1b–d). Taken together, these results indicated that the docking sites of exon 4 and 9 cluster are not essential for maintaining the overall transcript and protein levels of the *Dscam1* gene.

### Deletion of the docking site of exon cluster 6 markedly decreased normal exon 6 inclusion

When we used a similar strategy to remove the intronic docking site of exon 6 cluster, which has been conserved for ~500 million years of evolution^23^(Fig. 1a), the DscamΔ6D^−/−^ mutant showed adult homozygous lethality. To elucidate how deleting the docking site affected the inclusion of variable exon 6, we isolated DscamΔ6D^−/−^ homozygous embryos with a GFP marker. Only ~16% of the RT-PCR product was equal to the normal variable exon 6 in DscamΔ6D^−/−^ embryos (Fig. 1e,f). This result was in marked contrast to the splicing patterns observed in DscamΔ4D^−/−^ and DscamΔ9D^−/−^ mutants, but was largely consistent with the marked decrease of normal exon 6 inclusion seen in cell transfection assay using *Drosophila melanogaster*/*Drosophila virilis Dscam1* mutant constructs^26^. In the latter case, deletion of the docking site of exon cluster 6 caused abundant exon 6 skipping. These differences are likely to be due to different experimental approaches. Further sequencing analysis indicated that a cryptic exon was located immediately downstream of the docking site (Fig. 1e, f). As shown in Figure 1e, the larger band than normal exon 6 inclusion corresponded to the abnormal product of cryptic exon splicing to variable exon 6, while the smaller abnormal band corresponded to the product of cryptic exon splicing to constitutive exon 7. When the docking site was deleted, the spliceosome could aberrantly select the immediately downstream cryptic splice sites. In this scenario, the conserved intronic docking site also acted to repress the downstream cryptic splice sites. In addition, deletion of the docking site caused partial exon 6 skipping (Fig. 1e,f). In these cases, splicing of the cryptic exon to either variable exon 6 or constitutive exon 7, or exon 6 skipping, would introduce in-frame stop codons, leading to marked reduction of the overall normal protein level. As silencing of endogenous *Dscam1* expression by RNA interference recapitulated the *Dscam1* “null” lethal phenotype^31^, we speculated that DscamΔ6D^−/−^ lethality may be largely attributable to a marked reduction of Dscam1 protein level. As DscamΔ6D^−/−^ mutants showed adult homozygous lethality, we focused on the viable DscamΔ4D^−/−^ and DscamΔ9D^−/−^ mutants for alternative splicing and phenotypic analyses.

### DscamΔ4D^−/−^ shows global alteration of variable exon 4 bias

To investigate the effects of docking site disruption on variable exon choice, we first examined the relative usage of exon 4 variants in the RT-PCR products containing exon 4 using high-throughput sequencing. Docking site deletion resulted in a striking and profound change in the frequency of most exon 4s (Fig. 2a,b). Interestingly, the use of exons close to constitutive exons 3 and 5 (exons 4.1, 4.12) significantly increased in DscamΔ4D^−/−^, while exons in the middle-back region of the exon 4 cluster (exons 4.5, 4.6, 4.8, 4.9 and 4.11) decreased remarkably in frequency (Fig. 2a). To explore how the deletion of the docking site affected the inclusion of exon 4 variants, we analyzed the potential base pairing between the docking site and the selector sequences. In contrast to a recent study^32^, our evolutionary conservation analysis revealed that the selector sequences of at least four exon 4 variants (exon 4.2, 4.3, 4.6, 4.8) could potentially pair with the docking site (Fig. 2c; Supplementary Fig. 2), except for the previously identified exon 4.11^27^. Importantly, this base pairing prediction was strongly supported by the observation that five more highly conserved selector sequences were located in the *Dscam1* variable exon ortholog of housefly, a member of the common section Schizophora with an estimated divergence time of ~80 million years ago (Fig. 2c-f). Phylogenetic analyses reveal that 12 exon 4s of housefly *Dscam1* are orthologous to those of *Drosophila* species, suggesting a common mechanism for exon 4 orthologs of housefly and *Drosophila Dscam1*. These data together with previous experimental evidences using disrupting and compensatory mutations in *Drosophila* and hymenopteran *Dscam1* ^27, 28^, showed that the docking site contributed to determination of exon 4 inclusion frequency through base pairing with the selector sequence.

**Figure 2.**
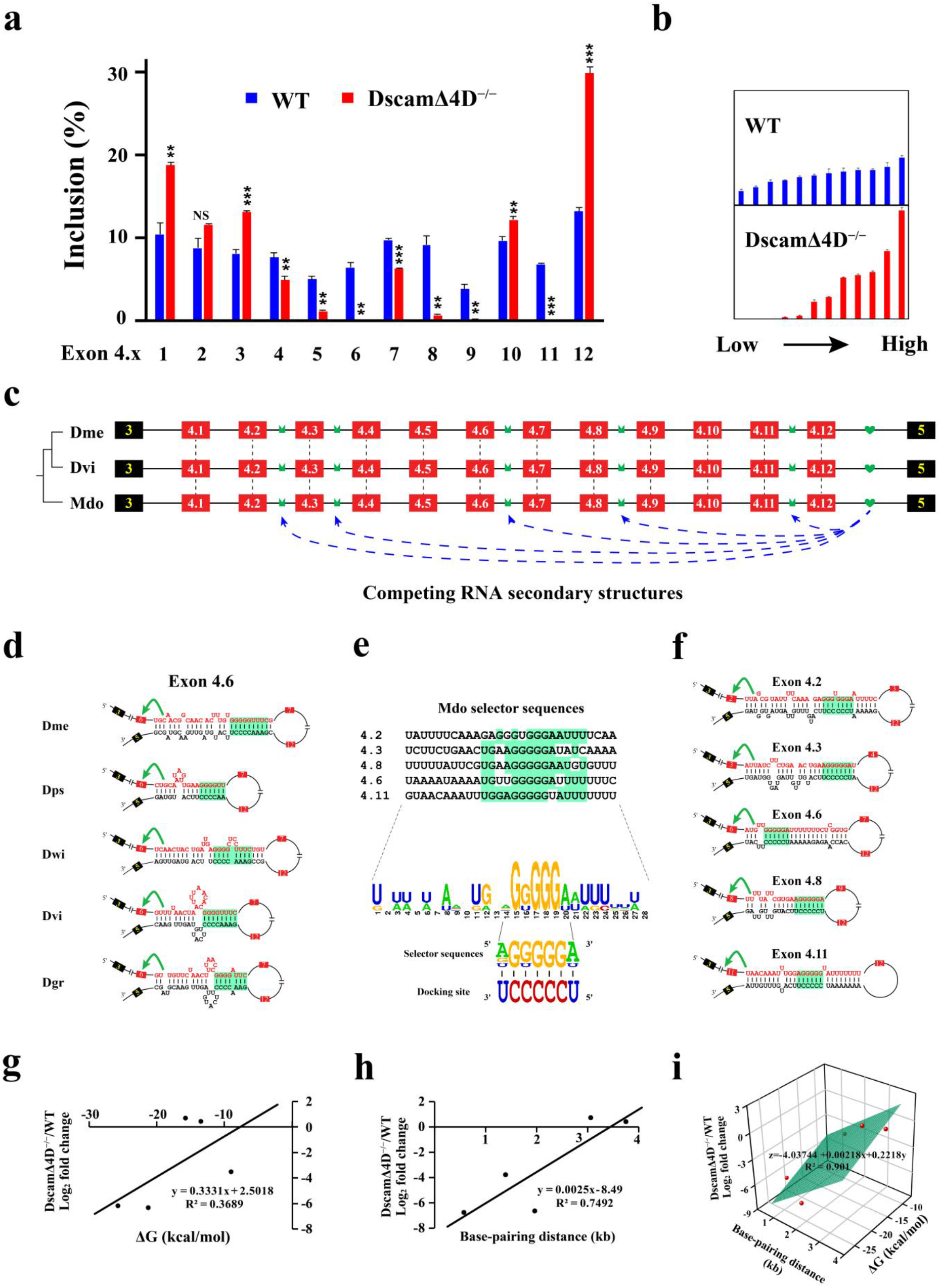
Deletion of docking site of exon cluster 4 globally altered isoform bias. (**a**) Deletion of the docking site dramatically impacted the choice of exon 4 variants; the inset shows comparisons of the relative frequency of the exon 4 inclusion between wild type and DscamΔ4D^−/−^ from minimal to maximum levels. Symbols used are the same as those in Fig. 1. The RT–PCR products from the wild type and DscamΔ4D^−/−^ animals were excised and sequenced using the multiplex sequencing method; all data are expressed as means ± SDs of three independent experiments. *P < 0.05; **P < 0.01; ***P < 0.001; NS, not significant (Student’s t-test, two-tailed). **(b)** Comparisons of the relative frequency of the exon 9 inclusion between wild type and DscamΔ4D^−/−^ from minimal to maximum levels. DscamΔ9D^−/−^ animals expressed a similar broad yet distinctive spectrum of exon 9 variants compared to the wild type. **(c)** Overview of variable exon 4s of *D. melanogaster* (Dme), *D. virilis* (Dvi), and *M. domestica* (Mdo) *Dscam1*. Phylogenetic analyses reveal that 12 exon 4s of housefly *Dscam1* are orthologous to those of *Drosophila* species (marked in dashed line), suggesting a common mechanism for exon 4 orthologs of housefly and *Drosophila Dscam1*. Our evolutionary conservation analyses revealed that selector sequences of at least four variable exon 4s (exon 4.2, 4.3, 4.6, 4.8) were potentially paired with the intronic docking site, except for previously identified exon 4.11 ^27^. **(d)** The base pairings between the docking site and selector sequence are conserved in exon 4.6 across *Drosophila* species. The base pairings between the docking site and selector sequence in exon 4.2, 4.3, 4.8 across *Drosophila* species were shown in Supplementary Fig. 2. The sequences that make up the core of the stem are highlighted in green. **(e)** The selector sequence consensus in *M. domestica*. The most frequent nucleotides in the central portion of the alignment are highlighted in green. The consensus nucleotides of the selector sequences were complementary to the docking site. These data clearly revealed one to many base pairings between the intronic docking site and selector sequence in *M. domestica*. (**f**) RNA secondary structures between the docking site and selector sequence in *M. domestica Dscam1*. (**g,h**) Analysis of correlations of the strength (g) or distance (h) of the base-pairing between the docking site and selector sequence with the fold change in the frequency of exon 4 inclusion. **(i)** Multiple linear regression analysis of exon 4 variants. This result revealed a much stronger correlation of the fold change in exon 4 inclusion with the combined effect of both the distance and strength of the docking site-selector base pairing than with either its strength or distance.

As expected, deleting the docking site almost abrogated the inclusion of exons 4.6, 4.8 and 4.11 because the base pairing interactions was disturbed in DscamΔ4D^−/−^ (Fig. 1a; Fig. 2d; Supplementary Fig. 2). Surprisingly, docking site deletion did not result in a substantial decrease, in fact an increase, in the inclusion frequency of exons 4.2 and 4.3 (Fig. 2a), which are located relatively far from the docking site. Considering the negative correlation of the looping distance between the docking site and selector sequence with variable exon inclusion^27^, we reasoned that the inhibitory effect of docking site deletion on exon 4.2 or 4.3 inclusion was gradually reduced as these two exons were far from the docking site (Fig. 2a,c). To address this, we performed a multiple linear regression analysis of the fold change in exon 4 inclusion against the strength and distance of the base pairing between the docking site and selector sequence. The analysis showed a much stronger correlation of the fold change in exon 4 inclusion with the combined effect of both the distance and strength of the docking site-selector base pairing (Fig. 2g-i). Taken together, these results indicate that the docking site deletion affected the inclusion frequency of exon 4 variants through disturbing the base pairing interaction in a proximity- and strength-mediated manner.

### Deletion of the docking site of exon cluster 9 globally changes variable exon 9 choice

Similarly, a global change in exon 9 usage was observed in DscamΔ9D^−/−^ flies. The spectrum of exon 9 variants in the heads of DscamΔ9D^−/−^ flies was similar to that of the wild type, but a marked change was observed in the relative usage of most exon 9 variants (Fig. 3a). Compared to wild type controls, 19 of 33 exon 9s were significantly increased, while five were significantly decreased, in DscamΔ9D^−/−^ flies. Interestingly, the relative usage of exons close to constitutive exon 10 (exons 9.30–9.33) decreased markedly. Similar changes in exon 9 usage were observed in various tissues of DscamΔ9D^−/−^, although the alterations were subject to developmental and tissue-specific regulation (Fig. 3b; Supplementary Fig. 3). These observations indicated that the docking site plays an important role in determining exon 9 inclusion frequencies in *Drosophila*.

**Figure 3.**
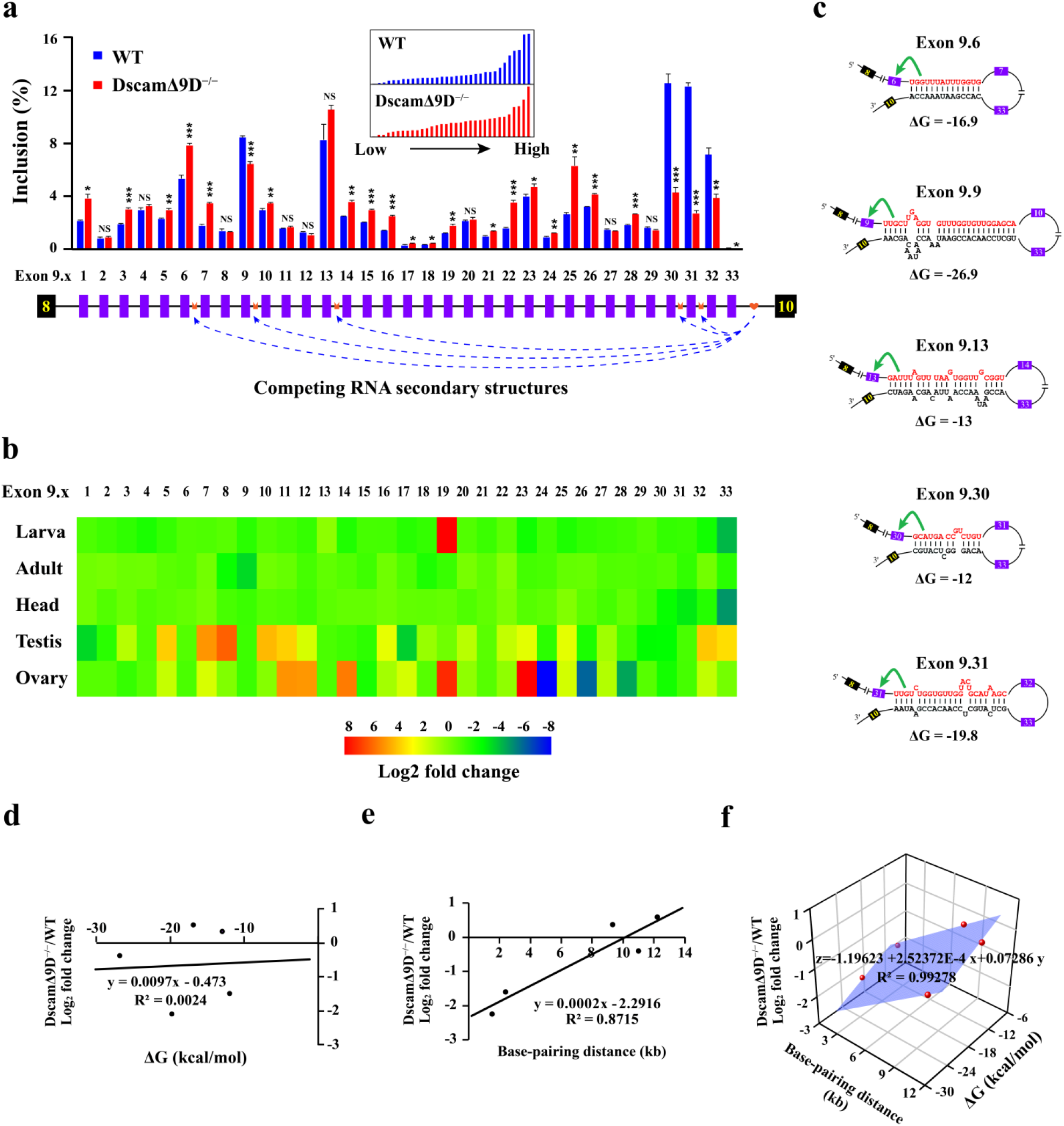
Deletion of docking site of exon cluster 9 globally altered isoform bias. (**a**) Deletion of the docking site dramatically altered the choice frequency of the exon 9 variant. Symbols used are the same as those in Fig. 1. The inset shows comparisons of the relative frequency of the exon 9 inclusion between wild type and DscamΔ9D^−/−^ from minimal to maximum levels. DscamΔ9D^−/−^ animals expressed a similar broad yet distinctive spectrum of exon 9 variants compared to the wild type. The RT–PCR products from the wild type and DscamΔ9D^−/−^ animals were excised and sequenced using the multiplex sequencing method; all data are expressed as means ± SDs of three independent experiments. *P < 0.05; **P < 0.01; ***P < 0.001; NS, not significant (Student’s t-test, two-tailed). (**b)** Heat map of changes in the frequency of the exon 9 inclusion in various developmental stages and tissues. The log_2_ fold change in each mutant construct compared to wild type is represented as the heat map. **(c)** RNA secondary structures between the docking site and selector sequence in exon 9 cluster of *D. melanogaster Dscam1*. RNA secondary structure between the docking site and selector sequence for exon 9.9 was previously described^27^. **(d,e)** The strength (d) or distance (e) of the base-pairing between the docking site and selector sequence is plotted as a function of the fold change in the frequency of the exon 9 variant. **(f)** Multiple linear regression analysis of exon 9 variants. The combined effect of both distance and strength of the predicted docking site-selector sequence interaction is plotted as a function of the fold change in the frequency of the exon 9 variant. The analysis revealed a much stronger correlation of the fold change in exon 9 inclusion with the combined effect of both the distance and strength of the docking site-selector base pairing than with either its strength or distance.

To determine how deletion of the docking site affected variable exon 9 selection, we analyzed the five most dominantly expressed variable exon 9s (exon 9.6, 9.9, 9.13, 9.30, 9.31). In contrast to a very recent study^33^, we found that their selector sequences were conserved across *Drosophila* species and were predicted to pair strongly with the docking site (Fig. 3c; Supplementary Fig. 4), except for the previously predicted exon 9.9 ^27^. As expected, deletion of the docking site led to marked reductions in exon 9.9, 9.30, and 9.31 inclusion, as base pairing interactions between the docking site and the selector sequences were disturbed (Fig. 1a; Supplementary Fig. 4). However, deletion of the docking site did not result in reduced inclusion of exon 9.6 or 9.13, and instead actually caused increased inclusion of these two exons in most tissues compared to wild type controls (Fig. 3a,b; Supplementary Fig. 3). Similar to exon 4 cluster, we reasoned that the inhibitory effect of docking site deletion on exon 9.6 or 9.13 inclusion was gradually reduced as these two exons were farther from the docking site than exon 9.30 or 9.31 (Fig. 3a). Notably, although exon 9.9 was located between exons 9.6 and 9.13, the strength of base pairing between the docking site and the selector sequence of exon 9.9 (ΔG −26.9 kcal/mol) was much stronger than that of exons 9.6 (ΔG −16.9 kcal/mol) and 9.13 (ΔG −13 kcal/mol) (Fig. 3c). This may explain why the docking site deletion markedly reduced inclusion of exon 9.9 but not exon 9.6, even though exon 9.9 resided farther from the docking site than exon 9.6. These observations together with disrupting and compensatory mutations on RNA secondary structures in exon 9 cluster of silkworm *Dscam1* ^28^, indicated that deletion of the docking site altered variable exon inclusion through perturbing base pairing-mediated regulation.

Similar to exon cluster 4, we observed a much stronger correlation of the fold change in exon 9 inclusion with the combined effect of both the strength and distance of the docking site-selector base pairing (Fig. 3d-f). A similar trend was observed in comparisons of two or more variables in correlation analyses of various tissues, although the correlation coefficients varied strikingly (Supplementary Fig. 5). Taken together with the results of exon 4 analysis, these observations indicated that deletion of the docking site in DscamΔ4D^−/−^ and DscamΔ9D^−/−^ altered variable exon inclusion through perturbing base pairing-mediated regulation in a proximity- and strength-mediated manner.

Consistent with previous studies^6, 21, 34^, comparative analyses did not reveal any significant difference between wild type and DscamΔ4D^−/−^ flies in the inclusion frequency of most alternative exons 6 or 9, showing largely independent splicing of exon clusters 4, 6, and 9. (Fig. 4a; Supplementary Fig. 6a,b). A similar conclusion was reached in analyses of inclusion frequency of alternative exons 4 and 6 in DscamΔ9D^−/−^ mutants (Fig. 4b; Supplementary Fig. 6c,d). In summary, we showed that deletion of the docking site perturbed base pairing-dependent regulation of splice isoform bias in a cluster-wide manner. Importantly, we successfully constructed DscamΔ4D^−/−^ and DscamΔ9D^−/−^ alleles encoding the same potential number of isoforms and overall expression level as wild type controls, albeit with globally changed exon variant bias (Fig. 4c). Therefore, these viable homozygous *Dscam1* mutants allowed us to test whether and to what extent altered isoform bias affected neuronal wiring.

**Figure 4.**
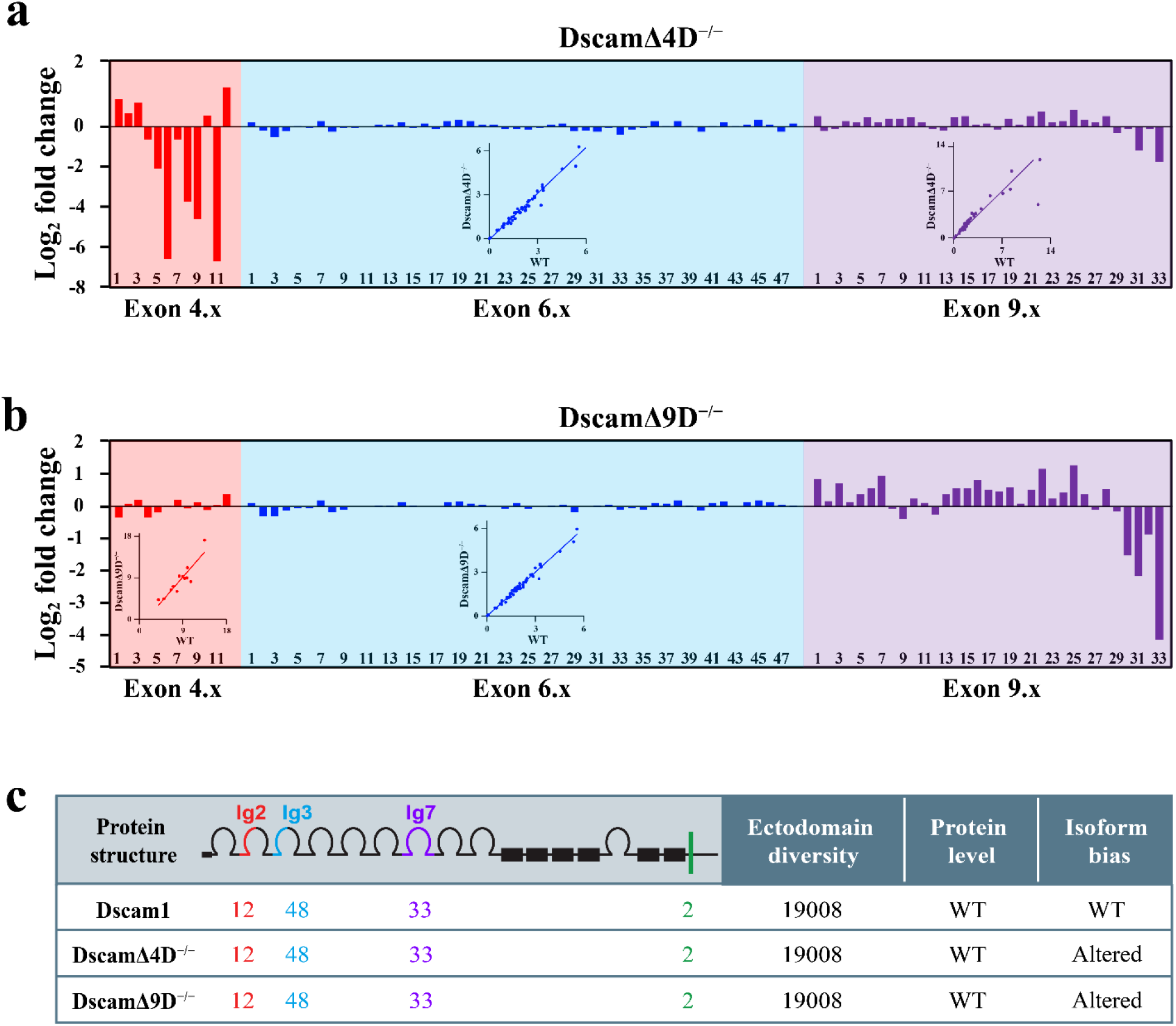
DscamΔ9D^−/−^ and DscamΔ4D^−/−^ show global alteration of isoform bias but with normal Dscam1 protein level. (**a**) The log_2_ fold change of the frequency of variable exon inclusion in wild type and DscamΔ4D^−/−^ was shown. The relative frequency of exon 4, 6, and 9 in DscamΔ4D^−/−^ is normalized to wild type control and presented as the log_2_ fold change. The correlation between wild type and DscamΔ4D^−/−^ is shown in the inset. This indicates that the docking site of the exon 4 cluster have almost not affected the inclusion of exon 6 and 9. (**b**) The log_2_ fold change of the frequency of variable exon inclusion in wild type and DscamΔ9D^−/−^ was shown. The correlation between wild type and DscamΔ9D^−/−^ is shown in the inset. This indicates that the docking site of the exon 9 cluster have almost not affected the inclusion of exon 4 and 6. (**c**) Characterization of the Dscam protein encoded by mutant *Dscam1* alleles, which expressed the same overall Dscam1 protein levels and same numbers of potential isoforms as the wild type control but had an altered isoform bias.

### Dscam1 isoform bias is dispensable for dendritic patterning

To investigate neuronal wiring in the mutant flies, we first examined whether altering Dscam1 isoform bias affect patterning dendritic arborization. The results did not reveal any significant differences between DscamΔ4D^−/−^ and wild type controls in self-crossing between sister branches of class I da (ddaE and vpda) neurons (Fig. 5a,b), suggesting that the marked alteration in isoform bias did not impair normal dendrite patterning. Identical results were obtained in DscamΔ9D^−/−^ flies (Fig. 5a,b). These results indicated that Dscam1 isoform bias was not required for dendrite morphogenesis of da neurons.

**Figure 5.**
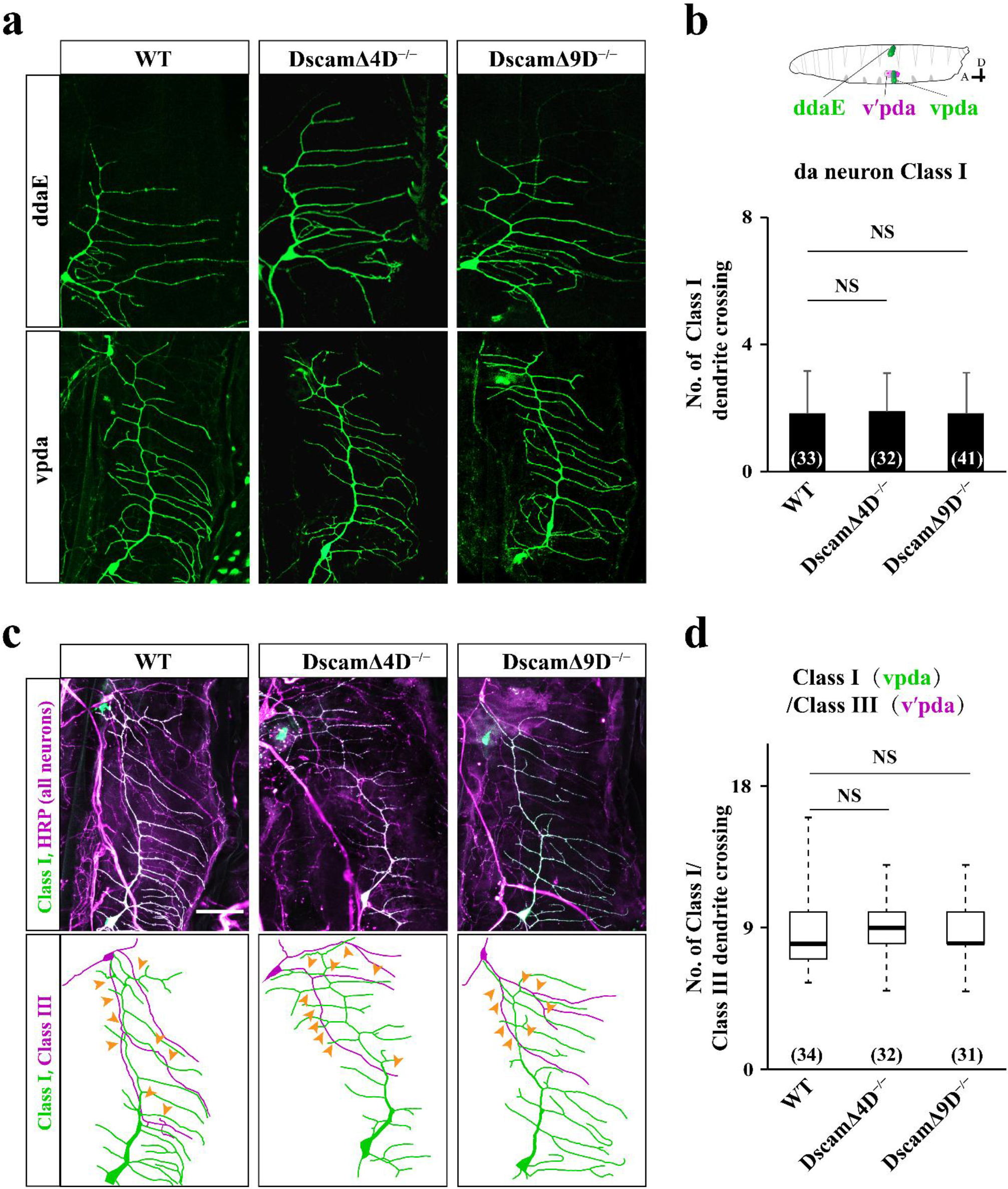
Dscam1 isoform bias is not required for the patterning of dendritic arborization neurons. (**a**) Representative images of class I da neuron dendrites from DscamΔ4D^−/−^, DscamΔ9D^−/−^, and wild type larval epidermis. Upper: ddaE neuron, lower: vpda neuron. (**b**) DscamΔ4D^−/−^ and DscamΔ9D^−/−^ da neurons show no significant differences in the number of dendrite crossings compared with wild type control. Numbers in parenthesis denote class I neurons analyzed for each mutant. Student’s t-tests, NS, not significant. (**c)** Representative images of dendritic arborization neuron dendrites in different *Dscam1* mutants. All neurons were visualized with anti-HRP antibody (magenta) while class I neurons were labelled with GFP (green; appears white because they overlap with magenta), and their corresponding color-coded tracings are shown below. Crossing between class I and III dendrites are shown with orange arrowheads. (**d)** No significant differences in the number of overlaps between class I and class III dendrites were identified between mutant and wild type. Numbers in parenthesis, class I/III pairs analyzed. Data in boxplot are represented as median (dark line), 25%–75% quantiles (box), and error bar (1.5× quartile range).

Previous studies indicated that Dscam1 diversity was pivotal for ensuring coexistence between dendrites from different da neurons^14, 15, 16, 18, 35^. To determine whether Dscam1 isoform bias was required for appropriate patterning of ensembles of dendrites, we examined the overlap between class I (vpda) and class III (v′pda) dendrites. Our observations indicated that class I dendrites showed considerable overlap with class III branches in third instar larvae of DscamΔ4D^−/−^ and DscamΔ9D^−/−^ mutants (Fig. 5c). Statistical analysis indicated no significant differences in the number of overlaps between class I and class III dendrites between mutants and wild type controls (Fig. 5d). Therefore, in contrast to the role of isoform number, Dscam1 isoform bias was dispensable for each da neuron to distinguish between self and non-self neurites.

### Altered isoform bias was associated with morphological defects in mushroom bodies in exon-specific manner

We next examined whether altering isoform bias influenced mushroom body development. To address this, the overall mushroom body morphology was visualized by staining with anti-FasII antibody (Fig. 6a). Compared to the completely normal phenotype of the wild type brains, up to 20% of the DscamΔ9D^−/−^ mutant brains exhibited mushroom body defects (Fig. 6b,c). In contrast to the *Dscam1* null phenotype, these mutant defects frequently occurred at either hemisphere of the entire mushroom body. These primary phenotypic defects included shortened and missing lobes, as well as β lobe fusion (Fig. 6b,c). In particular, mutant mushroom bodies frequently exhibited truncated lobes. In the few remaining samples, one mushroom body lobe was thinner than the other or showed misguidance (Fig. 6b,c). Notably, these mushroom body defects were largely different from those observed in mutant animals encoded to reduce diversity, in which the absence of lobes and thinner lobes are the predominate defects^18, 35^. In contrast, thinner lobes were rarely observed in DscamΔ9D^−/−^, which suggested that disruption of isoform bias may affect the mushroom body phenotype via mechanisms different from those that reduce isoform number.

**Figure 6.**
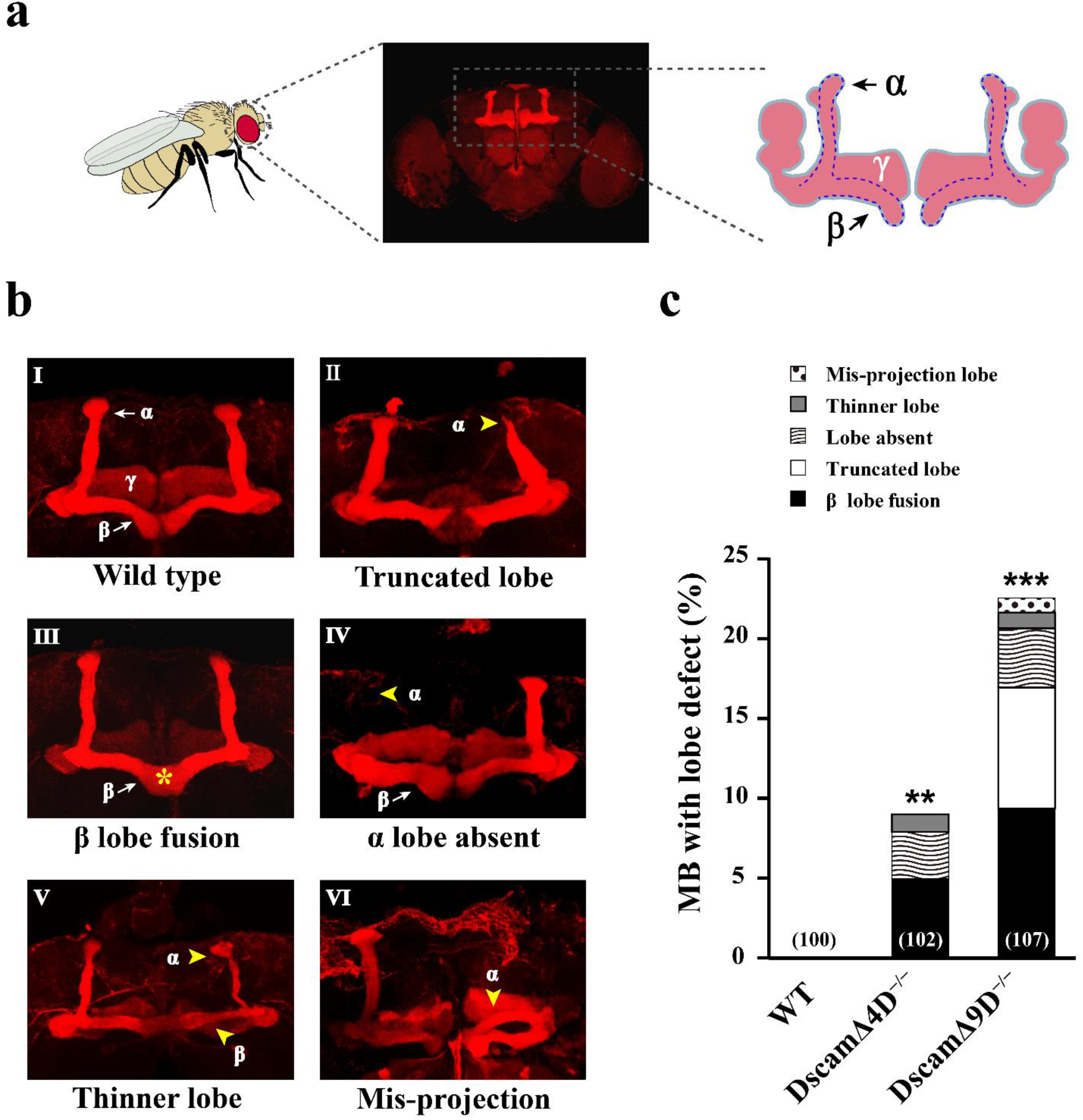
Disruption of Dscam1 isoform bias affected mushroom body morphology in exon-specific manner. (**a**) Schematics of mushroom body morphology of fly brain. (**b)** Representative images of mushroom body lobe morphology in mutant animals. Anti-FasII staining visualizing the α and β lobes of mushroom bodies in the adult brain of 3- to 7-day-old flies; yellow arrows indicate lobe defects. Scheme representing different mushroom body defects includes truncated lobes (II); β lobe fusion (III); lobe absent (IV); thinner lobes (V); mis-projection of lobes (VI). (**c**) Quantification of mushroom body defective phenotypes; numbers in parentheses represent the numbers of whole brains examined for each genotype. Data are expressed as mean±s.d. **P<0.01; ***P<0.001; (Student’s t-test, two-tail).

However, the DscamΔ4D^−/−^ animals exhibited a much less severe defective phenotype than DscamΔ9D^−/−^ animals (Fig. 6b,c). Except for defects in β lobe fusion, we have seldom observed mushroom body lobe defects (~2%, the number of brain hemispheres). This result is consistent with the previous observation that the morphology of mushroom bodies in Dscam1Δ4.1–4.3 or Dscam1Δ4.4–4.12 mutants (~2% defect) was indistinguishable from wild type^18^. Notably, both the number and frequency of exon 4 variants have been remarkably changed in these two deletion mutants^12, 18^. These data suggest that mushroom body morphology was not sensitive to disruption of exon 4 variant bias. As the number of Dscam isoforms and overall protein levels in DscamΔ9D^−/−^ and DscamΔ4D^−/−^ were identical to the wild type control, we concluded that the changes in isoform bias led to these gross mushroom body defects in the mutants. Taken together, these results indicated that Dscam1 isoform bias was required for precise mushroom body development in a variable exon-specific manner.

### Alteration of *Dscam1* exon 9 isoform bias is correlated with phenotypic defects

Having established the importance of *Dscam1* exon 9 isoform bias in mushroom body morphology, we next explored how the changes in exon 9 bias were associated with the defective phenotypes. To do this, we used CRISPR/Cas9 to delete the left and right portions of the docking site of exon cluster 9 from the *Dscam1* locus (Fig. 7a, b). We generated a series of knock-out mutants with varying degrees of deletions of the docking sequence in the exon 9 cluster (designated DscamΔ9D1–5^−/−^, Fig. 7a). This change was expected to have diverse effects on isoform bias by impacting docking site-selector base pairing interactions (Fig. 7b). Similar to the wild type controls, these mutants had normal overall *Dscam1* expression levels (Supplementary Fig. 7a–c), whereas the relative usage of the exon 9 variant was altered to different extents (Fig. 7d; Supplementary Fig. 7d). Thus, we created a series of mutant alleles exhibiting normal *Dscam1* expression levels but with altered isoform bias, which enabled us to investigate how varying the exon 9 isoform bias affects mushroom body morphology. As a result, these mutants exhibited different degrees of phenotypic defects (Fig. 7d). DscamΔ9D1^−/−^ and DscamΔ9D2^−/−^ showed modest changes in the relative frequency of exon 9 inclusion but no obvious phenotypic defects, whereas DscamΔ9D5^−/−^ exhibited more phenotypic defects, consistent with its marked changes in relative frequency of exon 9 inclusion (Fig. 7d). To determine the degree of correlation between altered isoform bias and phenotypic defects, we plotted the relative extent of isoform bias disruption against gross mushroom body defects. Statistical analysis revealed a striking linear correlation between the extent of disrupted isoform bias and mushroom body defects (Fig. 7d), which further confirmed the essential role of exon 9 isoform bias in normal mushroom body development.

**Figure 7.**
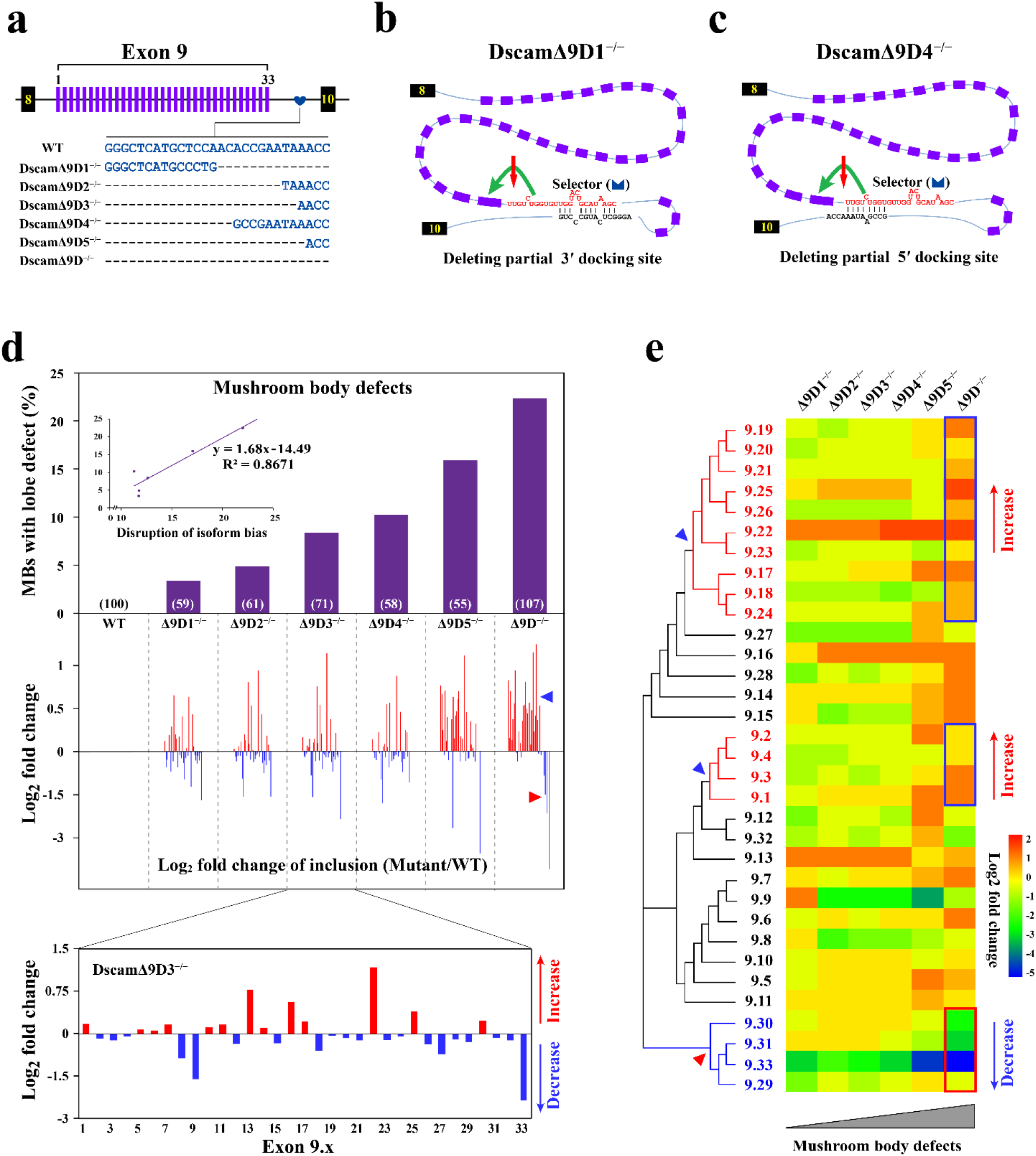
Mushroom body defects are correlated with the extent of disruption of isoform bias. (**a**) Schematic diagrams of docking site mutations in *D. melanogaster Dscam1* exon 9 cluster. We generated flies varying the degree of deletions of the docking site of the exon 9 cluster (designated as DscamΔ9D1–5^−/−^). (**b-c**) Schematic representation of RNA secondary structures altered by partially deleting the docking sites. The 3’ side of docking site was deleted (DscamΔ9D1^−/−^), or the 5’ side was deleted (DscamΔ9D4^−/−^). (**d)** The mushroom body defects are correlated with the extent of disruption of isoform bias. The log_2_ fold change of the frequency of exon 9 inclusion in wild type and mutants was shown below (from exon 9.1 to exon 9.33). The relative frequency of exon 9 inclusion in each mutant construct is normalized to wild type control and presented as the log_2_ fold change (from exon 9.1 to exon 9.33). The extent of disruption of isoform bias is estimated by total absolute value of the log_2_ fold change. Correlation of mushroom body defects with the extent of disruption of isoform bias is shown in the inset. Variable exon 9s with consecutive increases or decreases are shown as red and blue arrowheads. (**e**) Heat map of changes in the frequency of the exon 9 inclusion and their evolutionary relationship; the log_2_ fold change in each mutant construct compared to wild type is represented as the heat map. Variable exon 9s with consecutive increases or decreases in evolutionarily distinct clades (shown as red and blue arrowheads) are boxed.

Notably, there were no obvious phenotypic defects in some mutants (e.g., DscamΔ9D1^−/−^) despite substantial changes in their isoform biases (Fig. 7d). We concluded that the phenotypic defects were associated with a functional redundancy of the altered isoforms; indeed, the variable exon 9s tended to consecutively increase or decrease in DscamΔ9D^−/−^ and DscamΔ9D5^−/−^, while DscamΔ9D1^−/−^ showed the opposite trend (Fig. 7d). This was interesting because adjacent isoforms tend to be closely related and to have higher functional redundancy. In DscamΔ9D^−/−^, altering the isoform bias resulted in superimposed effects in evolutionarily distinct clades and thus more severe defects (Fig. 7e). In contrast, the deleterious effects of isoform bias alteration in DscamΔ9D1^−/−^ may have been antagonized by high redundancy of the within-clade isoforms. Indeed, we have seldom observed a striking correlation between the frequency or change of specific exon 9 inclusion and mushroom body defects (Supplementary Fig. 8, 9). These findings showed that the disruption of phylogenetically unique subsets of isoforms likely caused marked defects even if the change in a certain isoform generally led to subtle defects.

### Altered isoform bias causes axonal defects in neuroblasts and single-cell clones in a variable domain-specific manner

To gain insight into the mechanisms underlying the mushroom body defects, mosaic analysis with a repressible cell marker (MARCM) technique^36^ was used to assess *Dscam1*-induced axonal phenotypes at the level of the neuroblast or single-cell clone. Whereas wild type clones showed a bifurcated axon with dorsally and medially extending branches (Fig. 8a,b), approximately 32% of the DscamΔ9D^−/−^ mutant clones exhibited either a growth or guidance axonal defect, or a combination thereof (Fig. 8c–f′,g; Supplementary Fig. 10a–e′). The primary axonal defects included truncation and overextension of core axon branches, and missing, thinning, and/or thickening of α/β axons (Fig. 8c–f′,g; Supplementary Figs. 10a–e′; 11). These axonal defects largely resembled the mushroom body morphological defects detected by anti-FasII staining, whether they were observed in isogenic mutant brains, neuroblast clones, or single neuron clones (Fig. 8c–f′,g; Supplementary Figs. 10a–e′,f; 11). Conversely, the severity of the axonal defects observed in *Dscam1* mutant MARCM clones showed a strong correlation with that of mushroom body defects (Supplementary Fig. 11g–i). Considering that one mushroom body is derived from four indistinguishable cell lineages^37^, and the striking resemblance between the axonal and mushroom body defects observed in the DscamΔ9D^−/−^, we speculated that the defective mushroom body phenotypes were largely caused by axonal growth and guidance defects.

**Figure 8.**
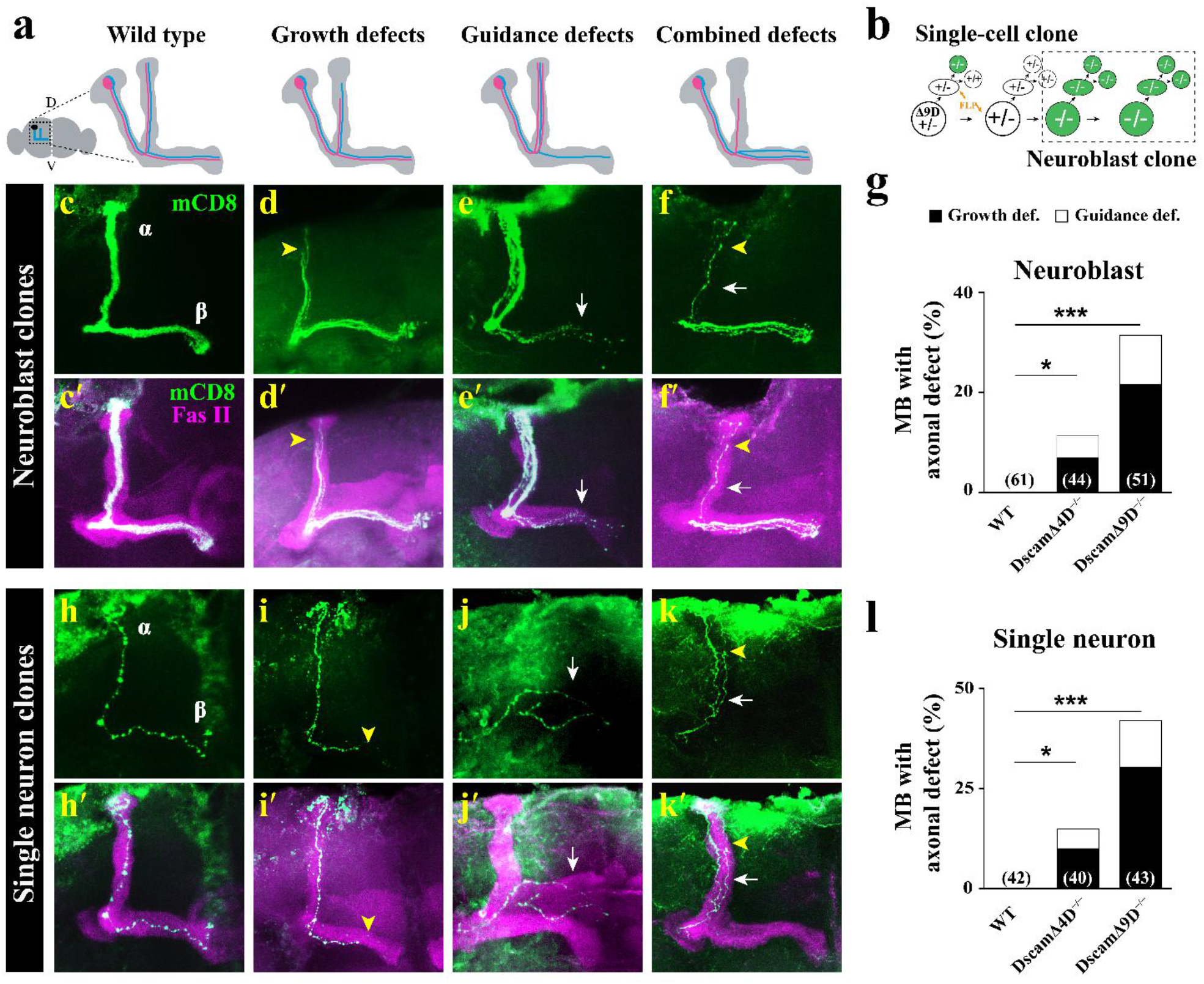
Phenotypic analysis of *Dscam1* mutant mushroom body axons at neuroblast and single-cell clones. (**a**) Schematic summary of axon growth and guidance defects in mutant mushroom body neurons. All drawings were of adult mushroom body of the brain hemisphere. A control neuroblast (NB) clone (WT) indicated the bifurcated axon with a dorsally and a medially running branch, while some DscamΔ9D^−/−^ mutant clones exhibited either axon growth defect or a guidance defect, or sometimes a combination of the growth and guidance defects. (**b**) Schematic illustration for selective labeling of NB clones or single neuron with MARCM strategy^36^. (**c**–**f**′) Wild type and DscamΔ9D^−/−^ mushroom body axons in NB clones. Compared with wild type NB clones (c, c′), mutant NB clones display axon growth (d, d′), and guidance (e, e′) defects, or sometimes a combination of them (f, f′). Yellow arrowhead indicated growth defect that axons did not reach the ends of α/β lobes in some mutant clones (Yellow arrowhead, d, d′). White arrowhead indicated guidance defect that axons in α lobe were thicker than in wild type control; and concomitantly, the axons in β lobe were thinner than normal (White arrowhead, e, e′). c–f were the same as c′–f′, respectively, showing FasII staining. (**g)** Quantification of mushroom body axon defects in NB clones. (**h**–**k′)** Wild type and DscamΔ9D^−/−^ mushroom body axons in single-cell clones. Unlike wild type control, DscamΔ9D^−/−^ exhibited either axon growth (i, i′), or guidance (j, j′) defects, or a combination of them (k, k′). h–k were the same as h′–k′, respectively, showing FasII staining. Yellow arrowhead indicated truncated axons and white arrowhead represented axonal misguidance. (**l**) Quantification of mushroom body axon defects in single-cell clones. Data are expressed as mean±s.d. from three independent experiments. *P<0.05; **P<0.01; ***P<0.001 (Student’s t-test, two-tail).

The axonal growth defects accounted for up to two thirds of the axonal defects, indicating that axon growth was most sensitive to disruption of Dscam1 isoform bias. The growth defects included truncation and overextension of mushroom body axon branches (Fig. 8c–f′). The former indicated that most processes did not reach the ends of α and β lobes in some mutant clones (Fig. 8d,d′), while the latter indicated that axon branches extended beyond the ends of α and β lobes (Supplementary Fig. 10b-c′). To gain further insight into the axonal growth defect, we examined single neuron clones homozygous for DscamΔ4D^−/−^ and DscamΔ9D^−/−^. We found that ~30% of homozygous DscamΔ9D^−/−^ single-neuron clones exhibited truncated or overextended axon branches, while the figure was less than 10% in DscamΔ4D^−/−^ single cell clones (Fig. 8l). Taken together, these phenotypic studies of single cell/neuroblast clones indicated that Dscam1 isoform bias is required to specify appropriate axon branch extension in a variable domain-specific manner.

We also found that axon guidance showed modest sensitivity to disruption of Dscam1 isoform bias. Guidance defects included absence, thinning, thickening, and misprojection of axons (Fig. 8e-f′; Supplementary Fig. 10d,e). When both α/β axon branches extended in the same lobe in these mutant clones, thickening of one lobe tended to be concomitantly paralleled with a thinning or lack of the other lobe (Fig. 8e-f′; Supplementary Fig. 10d,e). Single cell clone analysis revealed that the primary defect resulted from the failure of the normally divergent segregation of sister branches into the dorsal or medial lobe (Fig. 8j-k′). In addition, we observed supernumerary branches or an unusual single-branch phenotype without axon bifurcation at the peduncle end in some DscamΔ9D^−/−^ single cell clones, albeit at low frequency (Supplementary Fig. 10g-h′). Taken together, these phenotypic studies indicated that Dscam1 isoform bias is required for both normal growth and guidance of mushroom body axons.

The phenotypes of homozygous DscamΔ9D^−/−^ isogenic mutant brains showed lower penetrance than those of homozygous mutant neuroblasts and single neuron clones (26%, 32% versus 41%, Fig. 8g,l; Supplementary Fig. 11f). We speculated that disruption of isoform bias had cell non-autonomous effects on axon guidance and growth as in the *Dscam1* null mutant^9^. If each neuron independently determined its pathway to form axonal branches, we would expect to see a mixed pattern of wild type and mutant trajectories in neuroblast clones. Instead, all mushroom body axons in the DscamΔ9D^−/−^ neuroblast clones showed relatively similar types of guidance and growth defects. In these mutant neuroblast clones, all of the axons projected either dorsally or medially, rather than in divergent directions (Fig. 8e, j), or exhibited an either dorsal or medial single truncated lobe rather than truncation of both axonal lobes (Fig. 8d, i). The non-cell autonomous effects suggested that wild type axons may somehow dictate the projections of *Dscam1* mutant axons in mosaic mushroom bodies, and vice versa. Together, these observations indicated both cell autonomous and non-autonomous roles of Dscam1 isoform bias in regulating axon guidance and growth.

### Perturbation of Dscam1 isoform bias did not affect self-avoidance of axonal branches

Finally, we examined how disruption of Dscam1 isoform bias affected mushroom body axonal phenotypes. Two general scenarios may shed light on this issue. The first possibility is that disrupted isoform bias may alter the self-avoidance of axonal branches from the same neuron in which one sister branch is segregated into the dorsal lobe and the other into the medial lobe with high fidelity^35^. If this were true, we would expect alteration of isoform bias to cause fascicles and crossing between sister branches from the same neuron. The second possibility is that disruption of isoform bias influences Dscam1-mediated non-repulsive signaling strength, which in turn modulates axon growth and branching as described in mechanosensory axons ^17^.

To distinguish between these scenarios, we examined whether altered Dscam1 isoform bias influenced the self-avoidance of sister branches. Approximately 20% of DscamΔ9D^−/−^ axons were not reliably segregated into the dorsal and medial lobes compared to the completely normal projection pattern in the wild type axons (Fig. 9a, b). However, fascicles and crossing between sister branches were not observed in DscamΔ9D^−/−^ neurons, indicating that disrupted Dscam1 isoform bias did not affect the self-avoidance of sister branches from the same neuron. In contrast, sister branches in *Dscam1*^null^ neurons frequently crossed or formed fascicles^9, 11, 18, 35^(Fig. 9a,b). In addition, in contrast to single isoform mutants that exhibited homophilic interactions between sibling neurites from different neurons^14, 15, 16, 18, 35^, non-self branches from DscamΔ9D^−/−^ neurons should not engage in homophilic interactions that lead to self-avoidance because they encode the same potential repertoire of Dscam1 isoforms as wild type controls. Thus, the abnormal projections observed in DscamΔ9D^−/−^ mutants should not be attributed to self-avoidance between sister branches from the same neuron, or to self-avoidance between non-self branches from interneurons (tilling). Moreover, the growth defects (i.e., truncation) in the axon branches in DscamΔ9D^−/−^ mutants cannot be explained by the self-avoidance mechanism. These observations indicate that Dscam1 isoform bias mediates the patterning of mushroom body axonal wiring through influencing non-repulsive signaling.

**Figure 9.**
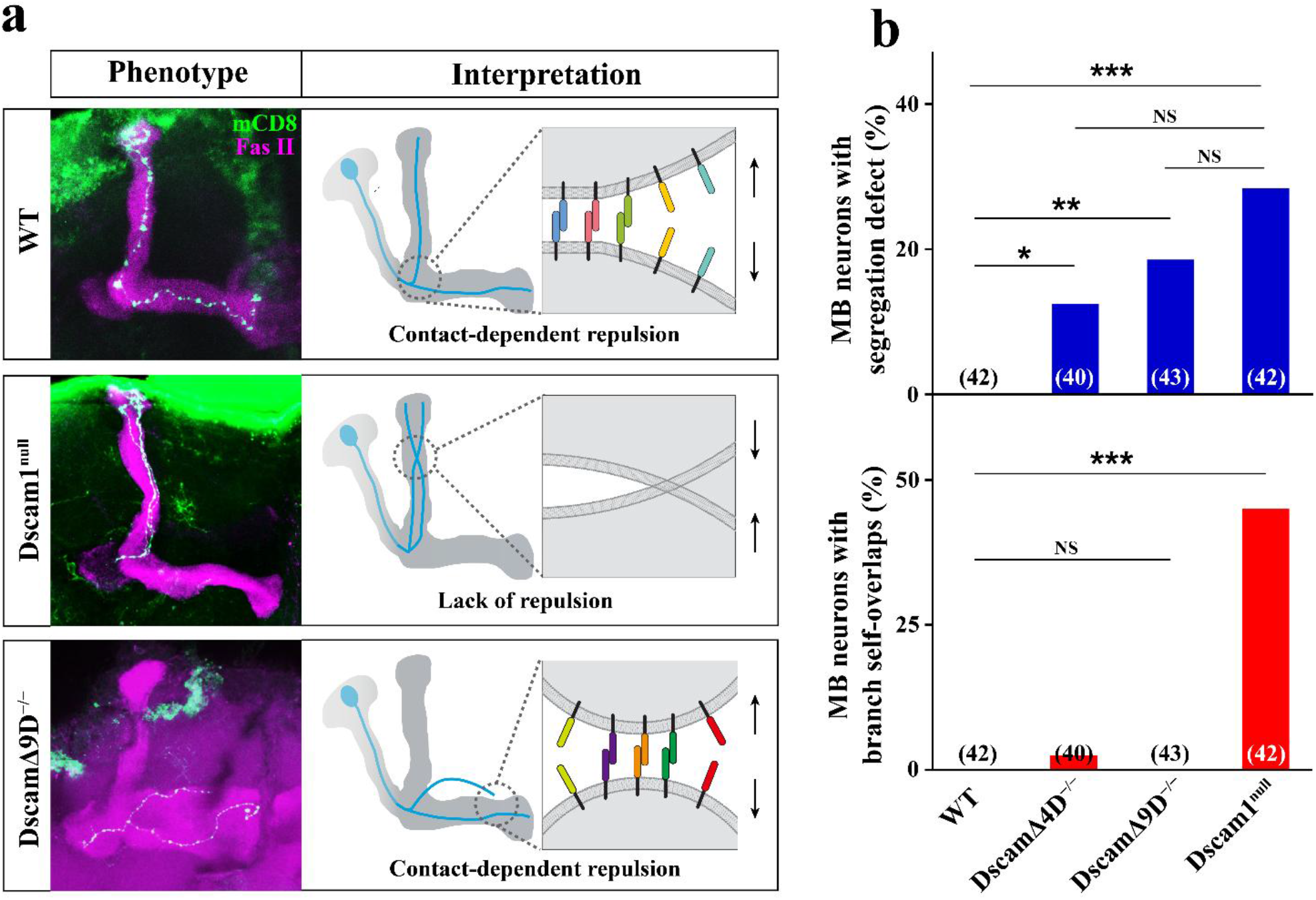
Disruption of Dscam1 isoform bias causes abnormal branch segregation but has no effects on the self-avoidance of sister branches. (**a**) Schematic of the effects of altered isoform bias on the self-avoidance of sister branches; representative mushroom body neurons are shown (blue). In the upper panel, the branches of wild type neurons exhibit little self-crossing due to contact-dependent repulsion mediated by Dscam1 isoforms (colored bars). By contrast, in the middle panel, the branches of *Dscam1*^null^ neurons frequently cross or form fascicles due to the loss of Dscam1-mediated repulsion. In the lower panel, DscamΔ9D^−/−^ animals exhibit a relative lack of self-crossing between abnormally segregated sister branches due to contact-dependent repulsion mediated by Dscam1 isoforms (colored bars). (**b**) Quantification of mushroom body axons with abnormal branch segregation or self-crossing in single-cell clones. Disruption of Dscam1 isoform bias led to abnormal branch segregation but a relative lack of self-crossing between sister branches. These observations reveal that disruption of Dscam1 isoform bias did not affect the self-avoidance of sister branches at a single-cell level. Data are expressed as mean±s.d. *P<0.05; **P<0.01; ***P<0.001; NS, not significant (Student’s t-test, two-tail).

## Discussion

This study provided the first conclusive evidence that Dscam1 isoform bias is essential for precise axonal wiring. Rather than deleting the coding exon region as in all previous studies, we used CRISPR/Cas9 to delete the intronic regulatory sequence. Surprisingly, deletion of the intronic docking sites did not affect the overall inclusion of the exon 4 or 9 clusters. This led to the generation of a genetic system appropriate for determining the importance of Dscam1 isoform bias. In contrast to the number of Dscam1 isoforms, which is important for distinguishing between self and non-self neurites by conferring unique single cell identity, our findings suggested that isoform bias may mediate mushroom body axonal wiring through Dscam1 non-repulsive signaling.

### Mechanistic implications for *Dscam* alternative splicing regulation

The results presented here indicated that deletion of the intronic docking site perturbed base pairing-mediated regulation of variable exon inclusion, albeit with strikingly different patterns of splicing change among Dscam1 exon clusters 4, 6, and 9. DscamΔD6^−/−^ mutants lacking the docking site of exon cluster 6 show adult lethality, commensurate with the marked decrease of normal exon 6 inclusions. In contrast, DscamΔ9D^−/−^ and DscamΔ4D^−/−^ were viable and fertile and showed normal overall inclusion of the exon 4 or 9 clusters but global alteration of isoform bias. These observations suggest that deletion of the docking site did not destroy the integrity of mutually exclusive splicing in the exon 4 or 9 cluster. These discrepancies suggest differences in base pairing-mediated splicing regulation between *Dscam1* exons 4 and 9 versus the exon 6 cluster. We proposed that other mechanisms may act to compensate for the inhibitory effects on exon inclusion caused by docking site disruption in the exon 4 or 9 clusters. Recently, we found that upstream and downstream intronic RNA secondary structures evolved to regulate the inclusion of variable exons in hymenopteran *Dscam* exon 4 and lepidopteran and hymenopteran exon 9 clusters^28^. Considering the evolutionary conservation and similarity of changes in exon inclusion between hymenopteran and fly *Dscam1* upon deletion of the docking site, we speculated that mutually exclusive splicing in the exon 4 or 9 cluster may be regulated by dual docking site-selector base pairing interaction in *Drosophila*.

Importantly, we showed that deletion of the intronic docking sites led to long-range alteration of exon variants, but with normal protein levels, in *Dscam* exon 4 and 9 clusters. Such alterations in variant ratio could be further mediated by fine-tuning the extent of the docking site mutation (Fig. 7). These results not only increased our understanding of the mechanism of *Dscam* complex alternative splicing, but also provided a useful system to study the importance of isoform bias in the nervous system.

### Role of *Dscam1* splicing bias in neuronal wiring in exon-specific manner

Previous genetic studies focused on *Dscam1* itself or its alternative exons, and therefore we were unable to determine whether the phenotypic defects were caused by alterations of isoform number or bias or changes in overall protein level. Rather than disrupting the exon region, as in previous studies^3, 9, 11, 13, 17, 29, 30^, we targeted disruption of the intronic regulatory element with CRISPR/Cas9 to generate mutant flies with normal overall protein levels and diversities, but with long-range changes in isoform bias. These mutants showed remarkable mushroom body and axonal defects, emphasizing that disruption of isoform bias may explain these phenotypic defects. The importance of isoform bias was further underscored by the striking correlation between degree of disruption of isoform bias and the severity of phenotypic defects (Fig. 7). Taken together, these results showed that intronic element-mediated splicing bias is essential for normal neuronal development.

Furthermore, our results showed that Dscam1 isoform bias is relevant to specific biological functions in a variable domain-specific manner. Our genetic analysis indicated that the disruption of variable exon 9 bias (DscamΔ9D^−/−^) caused markedly more severe defects than did that of variable exon 4 bias (DscamΔ4D^−/−^), coincident with the more striking bias in alternative exon 9 usage. Studies of expression profiling indicated that different mRNAs are expressed without any clear preference for exon 4 or 6, albeit with a strikingly biased usage of exon 9 in different types of cells and tissues^6, 7, 10, 20, 21^. Coincidently, exon 9-encoding Ig7 domains are the binding sites of Dscam receptor with its Netrin ligand^38, 39, 40^. This raises the possibility that the alteration of Ig7 variant bias may disturb the neuronal wiring through interference with the balance between heterophilic (i.e., netrin) and homophilic binding of Dscam1. These data are suggestive of a unique requirement for highly specific biases of Ig7 variants, at least in nervous system development.

Importantly, the results presented here suggest a role of Dscam1 isoforms distinct from canonical self-avoidance in neural circuits. *Dscam1* undergoes stochastic yet highly biased splicing^8^. Splicing stochasticity is perfectly compatible with a canonical role in neural circuit assembly through self-avoidance^2^. In this case, it is the number of isoforms, and not their identity, which matters most as long as homophilic binding occurs^7, 11, 14, 18, 35^. In contrast, high splicing bias is incompatible with self-avoidance but is suitable for intercellular recognition and synaptic choice mediated by specific isoforms or subsets of Dscam1. The latter view was supported by our genetic evidence that a series of DscamΔ9Ds^−/−^ mutants with the same degree of diversity as wild type controls exhibited distinct phenotypic defects (Figs. 5; 6c). Furthermore, our mosaic analysis, indicating that alteration of isoform bias caused remarkable axonal defects but failed to affect self-avoidance of axonal branches (Fig. 9), further underscores the speculation that at least certain Dscam1 isoforms have been diversified to specific roles in neuronal development beyond self-avoidance. Taken together, we concluded that the roles of the Dscam1 isoforms in neurite self-avoidance require isoform diversity (number), whereas a distinct role essential for neuronal development (i.e., neuronal growth) is primarily mediated by certain subsets of isoforms.

### The possible mechanisms for the requirement of Dscam1 isoform bias

However, it remains unclear how Dscam1 isoform bias affects neuronal wiring. We considered several possible mechanisms by which Dscam1 isoform bias may affect neuronal wiring. First, disruption of Dscam1 isoform bias influences Dscam1-mediated signaling strength, which in turn modulates axon growth and branching^17^. In our study, DscamΔ9Ds^−/−^ mutants frequently exhibited truncated mushroom body lobes or axons (Figs. 5, 7). These phenotypic defects were analogous to the severely impaired axonal growth and branching of mechanosensory axons caused by reduced diversity^17^. As biochemical analyses indicated that the strength of isoform-specific homophilic interaction varies markedly among different exon variants (i.e., 19~58 for Ig2, 13~37 for Ig3; and 1~38 for Ig7)^5^, it is possible that disruption of isoform bias may alter the strength of these homophilic interactions in mutant flies. The resulting overactivation or oversuppression of Dscam1 signaling would thereby lead to nascent axon branch retraction or overextension. In this scenario, Dscam1 isoform bias is necessary to maintain normal Dscam1 signaling. However, this model is not sufficient to explain a variable domain-specific requirement for isoform bias.

The second mechanism involves a change in neuronal guidance, which in turn involves ligand–receptor interactions. Several studies have suggested that Dscam receptors may recognize distinct specific ligands. For example, the Ig7 domain of Dscam is necessary for netrin-1 binding^38, 39, 40^, and Dscam binds Slit-N via at least two binding sites in the first five Ig domains, with one site lying in the first three Ig domains^41^. Our observations that Dscam1 isoform bias functions in a variable domain-specific manner also supported this possibility. As alterations of Dscam1 isoform bias led to substantial guidance defects, such alterations of isoform bias may affect guidance through ligand–receptor interactions. Alternatively, disruption of Dscam1 isoform bias may alter recognition and adhesive interactions between axon branches from different neurons. As Dscam1 isoforms could initiate signaling underlying multiple functions via homophilic and heterophilic interactions^1, 41, 42, 43^, Dscam1 isoform bias may contribute to neuronal wiring via combinatorial mechanisms.

### *Drosophila* Dscam1 versus vertebrate Pcdhs

*Drosophila Dscam1* and mammalian clustered *Pcdhs* encode extraordinary numbers of protein isoforms from a single complex genomic locus. Like *Drosophila Dscam1*, clustered Pcdh isoforms mediate self-avoidance through isoform-specific homophilic interactions followed by contact-dependent repulsion^44, 45^. However, Pcdhs relies on stochastic promoter activity for single cell identity, for self-recognition and self/non-self discrimination. Although both families generate extraordinary numbers of protein isoforms via distinct mechanisms^46, 47^, both Dscam1 and Pcdh isoforms are expressed in a biased stochastic fashion^6, 10, 20, 21^. Given the striking molecular parallels and complementary phylogenetic distribution between Dscams and Pcdhs, these two different gene families were proposed to have similar roles in the nervous system^2, 48, 49^. Nevertheless, there is a fundamental discrepancy in that certain Pcdhs (i.e., the C-type isoforms) play specific roles in neural wiring, whereas each Dscam1 isoform seems to be functionally equivalent^18, 48, 50, 51^. The evidence presented here strongly suggests that certain isoforms, or specific subsets of isoforms, have unique functions at least in some processes involved in neural wiring, similar to mammalian Pcdhs. Further functional genetic studies specifically targeting certain isoforms are required to determine these unique roles.

## Methods

### Fly Stocks

All flies were cultured on standard cornmeal medium at 25°C unless exceptional condition was mentioned. For each cross, we used four to five virgin females and two males to obtain comparable levels of offspring density. The fly strain W^1118^ (stock: 5905, Bloomington) was used as the wild type (WT). {nos-Cas9}attP2, in which Cas9 is specifically expressed in the germline via the nanos promoter^52^, was used for embryo injection. if/Cyo was used as the balancer stock for mutants. The 221-GFP Gal4 stock was used to drive GFP expression in class I da neuron. Single-cell marking of mushroom body (MB) neuron was achieved using the MARCM stocks FRT42D and Elav-GAL4, hs-FLP, UAS-mCD8-GFP; FRT42D-GAL80/Cyo.

### Generating altered bias alleles

In order to delete the intronic docking site in the *Drosophila Dscam1* gene, we performed targeted deletion using the CRISPR/Cas9 system^52^. Besides, the *Dscam1* null mutant was also generated by the CRISPR/Cas9 system. Briefly, target sequences of two guide RNAs (sgRNA) (Supplemental Table 1) were designed to delete the docking site of exon cluster 4, 6, and 9 from the *D*. *melanogaster Dscam1* locus, respectively (Fig. 1a). Two sgRNA plasmids were mixed with equal amount. Then, the purified sgRNA plasmids were co-injected into the {nos-Cas9}attP2 embryos. Injections were performed by SIBCB, CAS (Core Facility of Drosophila Resource and Technology Shanghai Institute of Biochemistry and Cell Biology). After injection, each potentially mosaic fly (G0 fly) was set up in an individual cross with balancer strain if/Cyo. Next, the male offspring (G1 fly) was crossed with the same balancer virgins. These mutants were examined through extracting genomic DNA (using TaKaRa Universal Genomic DNA Extraction Kit) followed by sequencing of the PCR product individually. Finally, the mutant flies were backcrossed with W^1118^ for five generations to get rid of unwanted mutations and off-targets.

### RNA pairing predictions and phylogenetic analysis

The RNA secondary structures between the docking site and selector sequences were predicted using the Mfold program^53^. In order to decipher the evolutionary relationship among the variable exons of *D. melanogaster Dscam1*, we aligned the amino acid sequences of the variable exons using ClustalW2 (http://www.ebi.ac.uk/clustalw/index.html)^54^and generated the phylogenetic trees using MEGA^55^ with the unweighted pair group method with arithmetic mean (UPGMA). The log_2_-scale fold-change of each mutant compared to wild type were calculated and represented as heat map using Heatmap Illustrator.

### RT–PCR

Total RNA was extracted from 50 fly heads using Trizol (Invitrogen) according to manufacturer’s instruction. Total RNA was reverse transcribed with the specific primer (Supplemental Table 2) using SuperScript III system (Invitrogen). The RT product was amplified using Ex taq (TaKaRa) for variable exons 4, 6 and 9 clusters, respectively. Detailed PCR parameters include : denaturing at 95°C for 3 min, 35 cycles of denaturing at 95°C for 30 sec, annealing at 58°C for 30 sec, and extension at 72°C for 1 min, followed by extension at 72°C for 10 min. The RT–PCR products were examined by electrophoresis with 1.5% agarose gel.

### Quantitative PCR

Quantitative PCR (qPCR) was used to test whether overall *Dscam1* transcripts were altered in the mutant flies. Total RNA was isolated using the Trizol reagent (Invitrogen) according to the manufacturer’s protocol. RNAs were reverse-transcribed using oligo(dT)_20_ primer (Invitrogen). qPCR was performed using SYBR premix Ex Taq (TaKaRa) according to the manufacturer’s protocol. The qPCR primers were designed for exon 10 and 11 (Supplemental Table 2). *D. melanogaster Actin* was used as the control gene to normalize the qPCR results. The reaction was run on Bio-Rad myiq2 real time PCR detection system using the following protocol: 95°C for 3 min, followed by 30 cycles of annealing and amplification at 60°C for 1 min. As a final step, the products were heated up to 95°C with continuous fluorescence measurements to obtain the melting curves. The results were analyzed using the software Bio-Rad IQ5.

### Utilization assay of exon variants

The relative inclusion ratios of the alternative exon 4, 6, and 9 were determined by using high-throughput sequencing. The PCR primers were designed to amplify the alternative exon and its partial flanking constitutive exons (Supplemental Table 2). RT–PCR was performed under the following cycling conditions: denaturing at 94°C for 5 min, 25 cycles of denaturing at 94°C for 30 sec, annealing at 60°C for 1 min, and extension at 72°C for 1 min, and a final elongation step of 10 min at 72°C. The PCR products were examined by electrophoresis with 1.5% agarose gel. The target products were excised and gel purified using the Biospin Gel Extraction Kit (BioFlux). Purified PCR products were pooled on Illumina MiSeq platform (Illumina, San Diego, USA) according to the standard protocol by G-BIO.

### Western blot analyses

Protein samples were prepared using strong RIPA lysis buffer (CW-BIO) (50mM Tris pH 7.4, 150 mM NaCl, 1% Triton X-100, 1% sodium deoxycholate, 0.1% SDS, sodium orthovanadate, sodium fluoride, EDTA, leupeptin) containing the protease inhibitor PMSF (Beyotime). Equal amounts of total protein were separated by 12% SDS–PAGE gel, and were transferred onto nitrocellulose membranes. After blocking with 5% BSA (VWR LIFE SCIENCE) for 1 h at room temperature, the membranes were incubated with diluted primary antibodies to Dscam1 (ab43847, diluted 1:5000) or β-actin (ab8227, diluted 1:10,000) at 4°C overnight. After three washes with TBST, the membranes were incubated with secondary antibodies Goat Anti-Rabbit IgG (1:10,000, CW-BIO) at room temperature for 2 h. The immunoreactive bands of Dscam1 and actin were detected using the eECL Western Blot Kit (CW-BIO), and were measured by Tanon 5200.

### Immunostaining

All flies were between 3- to 7-day-old at the time of dissection. Adult brains were dissected in phosphate buffered saline (PBS). Then, the brains were fixed in 4% paraformaldehyde (diluted in PBST) at room temperature for 45 min, followed by three washing steps in PBS containing 0.1% Triton X-100 (PBST) for 20 min each. We blocked them in 5% BSA blocking solution (diluted in PBST) for 1 h at room temperature. After 20 min wash by PBST for three times, the brains were incubated for 48 h at 4°C with the primary antibodies diluted in PBST. The following antibodies were used: monoclonal antibody 1D4 (anti-FasII) (DSHB) diluted 1:2 in PBST, rabbit anti-GFP (Invitrogen) diluted 1:800 in PBST. Subsequently, brains were washed three times for 20 min each in PBST and were incubated for 4 h with the secondary antibodies at room temperature. The following antibodies were used: Alexa488-goat-anti-rabbit IgG or Alexa594-goat-anti-mouse IgG (Earthox) diluted 1:400 in PBST. After incubation with the secondary antibodies followed by standard washing step, the samples were mounted in prolong gold antifade reagent (Invitrogen). The immunofluorescence staining was imaged with a laser scanning confocal microscope LSM800 (Carl Zeiss).

We examined class I da neurons of third instar larvae as described previously^56^. The 221-GFP Gal4 stock was used to drive GFP expression in class I da neuron. We examined third instar larvae with GFP-labeled class I da neuron under the fluorescence microscope (Nikon SMZ18). The selected larvae were immersed in the PBS and were dissected along the dorsal or ventral midline. Larvae epidermis was fixed with 4% paraformaldehyde (diluted in PBST) for 1 h at room temperature. After rinsed for three times in PBST, the larvae epidermis was blocked in 5% BSA. Antibody (Cy3-conjugated Affinipure Goat Anti-Horseradish Peroxidase) was used at a concentration for 1:200 and was incubated overnight at 4°C. After standard washing step, samples were mounted in prolong gold antifade reagent (Invitrogen). The immunofluorescence staining was imaged with a laser scanning confocal microscope LSM800 (Carl Zeiss). Quantification of dendrite crossings between class I (vpda) and class III (v’pda) was represented in boxplots as median, quartiles Q1-Q3 (25%-75% quartiles), and error bar in 1.5 × quartile range.

### Induction and phenotypic analysis of MARCM clones

Single-cell clones and neuroblast (NB) clones of homozygous mutant α/β neurons were generated using MARCM techniques^36^. For mutant analysis, the DscamΔ4D^−/−^ and DscamΔ9D^−/−^ were crossed to FRT42D for chromosome recombination individually. FRT42D, DscamΔ4D^−/−^ or FRT42D, DscamΔ9D^−/−^ mutants were mated with Elav-GAL4, hs-FLP, UAS-mCD8-GFP; FRT42D-GAL80/Cyo to generate GFP expression clones. Then, mitotic recombination was induced via heat shock for 1 h in a water bath at 37°C. Immunostaining of adult brains was performed as described above.

### Statistical analysis

The results of the quantitative analyses were obtained from at least three biological replicates. The error bars were calculated from the average of three independent experiments. The significance of differences was determined by a two-tailed Student’s t-test, and *P<0.05, **P<0.01 and***P<0.001 were adopted to indicate statistical significance. Three-factor correlation was analyzed by multiple linear regressions. Statistical analysis was performed by multiple linear regression using Origin 2018 software (OriginLab). A multiple linear regression was employed to fit the data, taking the fold change in the frequency of exon variant as the dependent variable and the distance and strength of the base-pairing between the docking site and selector sequence as the two independent variables.

## Supplementary information

Supplementary Tables 1, 2

Supplementary Figs. 1–11

## Acknowledgments

We thank Haihuai He, and Yijun Meng for comments on the manuscript; and members of the Jin lab for suggestions and discussion during the course of this work. We are grateful to Yujie Fan, Ying Wang, Chun Hu for technical help. We thank Yan Li for the 221-GFP; Jianquan Ni for {nos-Cas9}attP2, U6a-sgRNA-short plasmid; Xiaohang Yang for if/Cyo, FRT42D and Elav-GAL4, hs-FLP, UAS-mCD8-GFP; FRT42D-GAL80/Cyo. This work was supported by research grants from the National Natural Science Foundation of China (31430050, 31630089, 91740104).

## Author contributions

YJ conceived of this project. WH, HD, JZ, FZ, YW, and SC designed and performed the experiments; WH, HD, JZ, FZ performed generating altered bias alleles; SC performed generation of Dscam1^null^; WH, BX, YS and FZ conducted the analysis of isoform expression; WH, HD, JZ, WY and YW performed phenotype analysis. YJ, FS, ZG and JH analyzed the data; YJ and WH wrote the manuscript; all authors discussed the results and commented on the manuscript.

