## Supplemental Material for "Intron-targeted mutagenesis reveals roles for *Dscam1* RNA pairing-mediated splicing bias in neuronal wiring"

### Supplementary information

Supplementary Tables 1, 2

Supplementary Figs. 1–11

| Table S1 Specific primers used for the sgRNA | | |
|---|---|---|
| **Primer** | **5'-3' sequence** | **Assay** |
| **Ds-4-sg-F1** | TTCGGGCAAGTCAGGAGCGACCCA | sgRNA for DscamΔ4D$^{-/-}$ |
| **Ds-4-sg-R1** | AAACTGGGTCGCTCCTGACTTGCC | sgRNA for DscamΔ4D$^{-/-}$ |
| **Ds-4-sg-F2** | TTCGCATATTGTTACGTGGTCGAT | sgRNA for DscamΔ4D$^{-/-}$ |
| **Ds-4-sg-R2** | AAACATCGACCACGTAACAATATG | sgRNA for DscamΔ4D$^{-/-}$ |
| **Ds-6-sg-F1** | TTCGAGTACCCTATCCCAACATTC | sgRNA for DscamΔ6D$^{-/-}$ |
| **Ds-6-sg-R1** | AAACGAATGTTGGGATAGGGTACT | sgRNA for DscamΔ6D$^{-/-}$ |
| **Ds-6-sg-F2** | TTCGACCTGGCGATTCAGTTAACT | sgRNA for DscamΔ6D$^{-/-}$ |
| **Ds-6-sg-R2** | AAACAGTTAACTGAATCGCCAGGT | sgRNA for DscamΔ6D$^{-/-}$ |
| **Ds-9-sg-F3** | TTCGGCTGATACGCAGTGGGACCT | sgRNA for DscamΔ9D$^{-/-}$ |
| **Ds-9-sg-R3** | AAACAGGTCCCACTGCGTATCAGC | sgRNA for DscamΔ9D$^{-/-}$ |
| **Ds-9-sg-F4** | TTCGGAGTCAGTCTCGTTTGGTGG | sgRNA for DscamΔ9D$^{-/-}$ and DscamΔ9D1$^{-/-}$ |
| **Ds-9-sg-R4** | AAACCCACCAAACGAGACTGACTC | sgRNA for DscamΔ9D$^{-/-}$ and DscamΔ9D1$^{-/-}$ |
| **Ds-9-sg-F5** | TTCGGTTATTGGTTTATTCGGTGT | sgRNA for DscamΔ9D2$^{-/-}$,DscamΔ9D3$^{-/-}$,DscamΔ9D4$^{-/-}$,DscamΔ9D5$^{-/-}$ |
| **Ds-9-sg-R5** | AAACACACCGAATAAACCAATAAC | sgRNA for DscamΔ9D2$^{-/-}$,DscamΔ9D3$^{-/-}$,DscamΔ9D4$^{-/-}$,DscamΔ9D5$^{-/-}$ |
| **Ds-9-sg-F6** | TTCGTGTAAAATACAAGTTCACAG | sgRNA for DscamΔ9D2$^{-/-}$,DscamΔ9D3$^{-/-}$,DscamΔ9D4$^{-/-}$,DscamΔ9D5$^{-/-}$ |
| **Ds-9-sg-R6** | AAACCTGTGAACTTGTATTTTACA | sgRNA for DscamΔ9D2$^{-/-}$,DscamΔ9D3$^{-/-}$,DscamΔ9D4$^{-/-}$,DscamΔ9D5$^{-/-}$ |
| **Ds-9-sg-F7** | TTCGCTTGCTGTTATTGGTTTATT | sgRNA for DscamΔ9D2$^{-/-}$,DscamΔ9D3$^{-/-}$,DscamΔ9D4$^{-/-}$,DscamΔ9D5$^{-/-}$ |
| **Ds-9-sg-R7** | AAACAATAAACCAATAACAGCAAG | sgRNA for DscamΔ9D2$^{-/-}$,DscamΔ9D3$^{-/-}$,DscamΔ9D4$^{-/-}$,DscamΔ9D5$^{-/-}$ |

| Table S2 Specific primers used for RT–PCR and PCR analyses | | |
|---|---|---|
| **Primer** | **5'-3' sequence** | **Assay** |
| **Ds-4-exa-F** | ATATCGCGGAAAGCGGTAAATTTACAG | Examine DscamΔ4D$^{-/-}$ |
| **Ds-4-exa-R** | CTATACACCAACAATGCCCAAGAGTTTGTA | Examine DscamΔ4D$^{-/-}$ |
| **Ds-6-exa-F** | CCGAAAAGAATTGTTGCTGTAG | Examine DscamΔ6D$^{-/-}$ |
| **Ds-6-exa-R** | ATTGGGAAACTTGGGCGAAACTCTG | Examine DscamΔ6D$^{-/-}$ |
| **Ds-9-exa-F** | TCTCTGAATCTATCCATATTCCACATCCGT | Examine DscamΔ9D$^{-/-}$ |
| **Ds-9-exa-R** | GTTTCCTTGCCAAAGGACTTCGCTTTGTGC | Examine DscamΔ9D$^{-/-}$ |
| **Ds-3-F** | TGGATCAGGAGCGACGGTAC | RTPCR |
| **Ds-5-R** | CTCCAGAGGGCAATACCAGG | RTPCR |
| **Ds-5-F** | GCTACCAGTGCCGAACCAAACATC | RTPCR |
| **Ds-7-R** | AGTCTCAACGCTTTCGCCTCCCAC | RTPCR |
| **Ds-8-F** | ACTTGCGTTGCCAAGAATCAGGAAG | RTPCR |
| **Ds-10-R** | GCCTTATCGGTGGGCTCGAGGATCC | RTPCR |
| **Ac-F-q** | TGTGGATATCCGTAAGGATC | QPCR |
| **Ac-R-q** | ACAGCGAAGCCAGGATGGAG | QPCR |
| **Ds-10-F-q** | TAAGGCCTTCGCCCAGGGATCC | QPCR |
| **Ds-11-R-q** | TCTCCGGGGGTGTCGCCAACT | QPCR |
| **Ds-3-F-seq** | TCGTCGGCAGCGTCAGATGTGTATAAGAGACAGTGGATCAGGAGCGACGGTAC | Sequencing |
| **Ds-5-R-seq** | GTCTCGTGGGCTCGGAGATGTGTATAAGAGACAGCTCCAGAGGGCAATACCAGG | Sequencing |
| **Ds-5-F-seq** | TCGTCGGCAGCGTCAGATGTGTATAAGAGACAGGCTACCAGTGCCGAACCAAACATC | Sequencing |
| **Ds-7-R-seq** | GTCTCGTGGGCTCGGAGATGTGTATAAGAGACAGAGTCTCAACGCTTTCGCCTCCCAC | Sequencing |
| **Ds-8-F-seq** | TCGTCGGCAGCGTCAGATGTGTATAAGAGACAGACTTGCGTTGCCAAGAATCAGGAAG | Sequencing |
| **Ds-10-R-seq** | GTCTCGTGGGCTCGGAGATGTGTATAAGAGACAGGCCTTATCGGTGGGCTCGAGGATCC | Sequencing |
| **Ds-10-RT** | GGTTTGGGGAAGCCATCAGCCTT | RT |

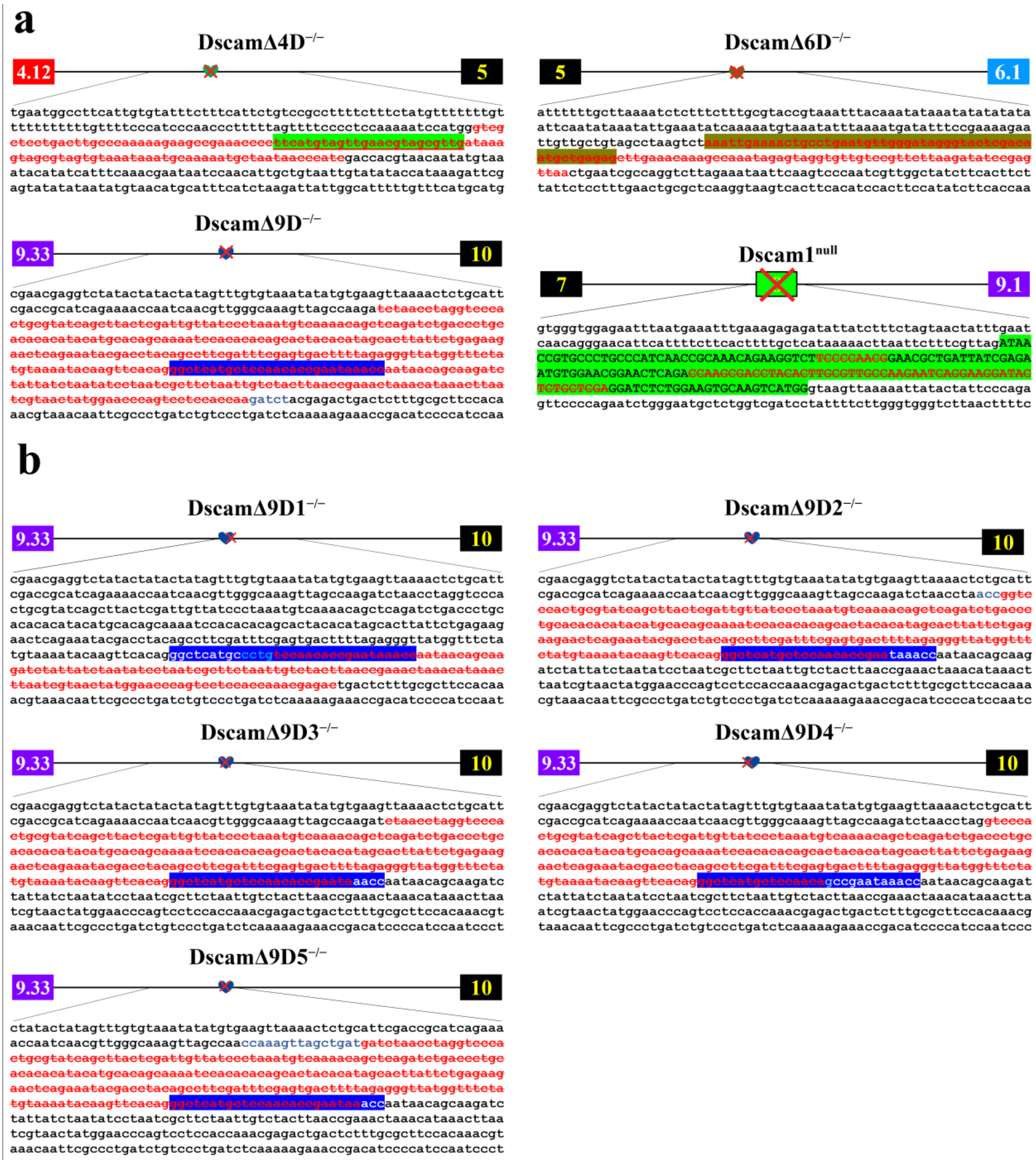


**Figure S1. Schematic diagrams and deleted sequences of fly *Dscam1* mutant alleles.** (a) Schematic diagrams and deleted sequences of DscamΔ4D$^{-/-}$, DscamΔ6D$^{-/-}$, and DscamΔ9D$^{-/-}$. The docking sequences (marked by hearts) of the exon 4, 6, and 9 clusters from the *D. melanogaster Dscam1* locus were deleted using the CRISPR/Cas9 system (designated as DscamΔ4D$^{-/-}$, DscamΔ6D$^{-/-}$, and DscamΔ9D$^{-/-}$). Correspondingly, we deleted the middle region of the constitutive exon 8 from *D. melanogaster Dscam1* locus (designated as *Dscam1*$^{null}$), which resulted in frameshift mutation (marked by a red cross). The deleted sequences are shown in red. (**b**) Schematic diagrams and deleted sequences of DscamΔ9D1–5$^{-/-}$. We used the CRISPR/Cas9 system to generate mutant flies varying the degree of deletions of the docking sequence of the exon 9 cluster (designated as DscamΔ9D1–5$^{-/-}$).

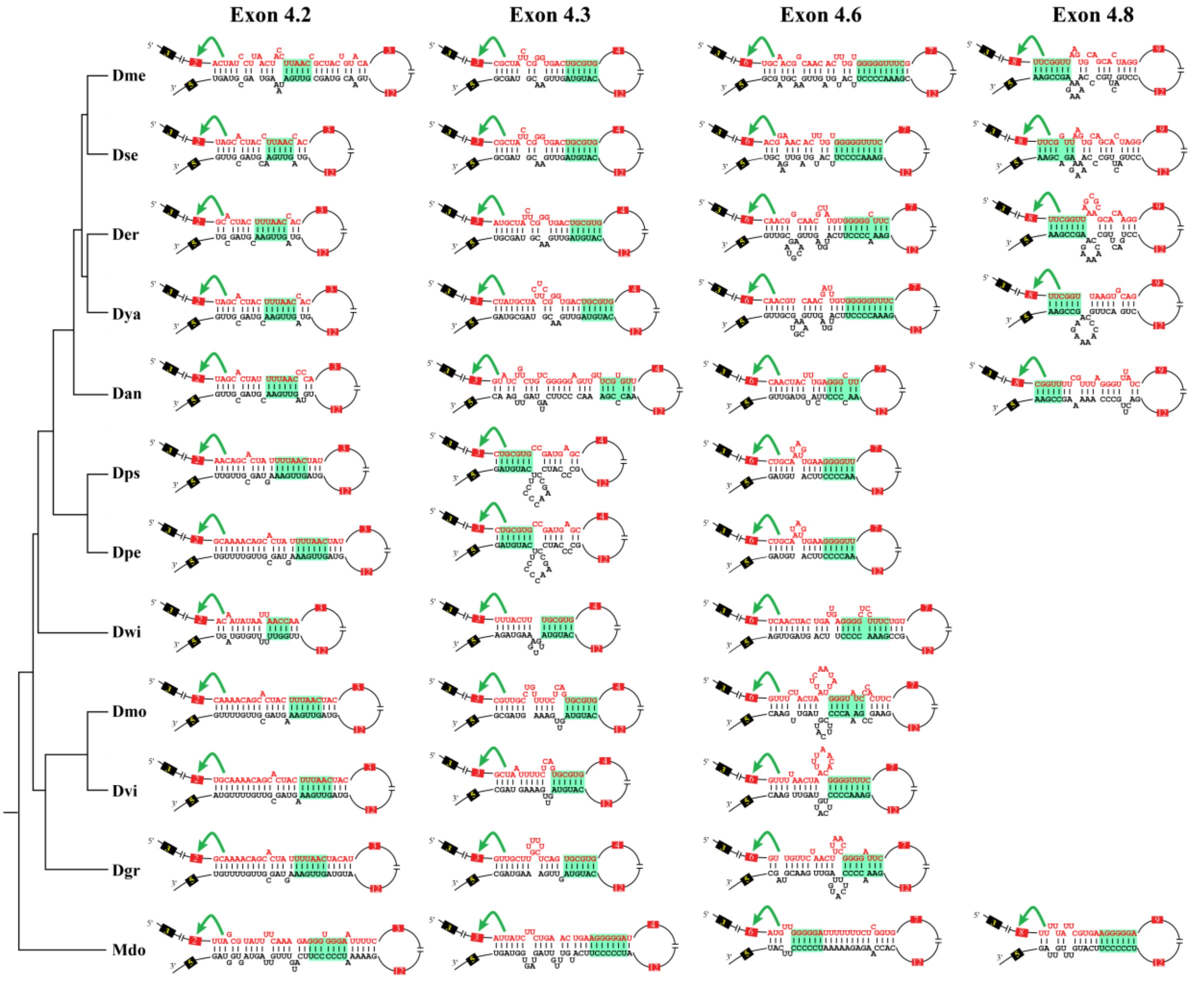


**Figure S2. RNA secondary structures between the docking site and selector sequences are conserved across *Drosophila* and housefly.** The base pairings between the docking site and selector sequence are conserved in exon 4.2, 4.3, 4.6, 4.8, 4.11 across *Drosophila* species and housefly. Those *Drosophila* species include *D. melanogaster* (Dme), *D. simulans* (Dsi), *D. sechellia* (Dse), *D. yakuba* (Dya), *D. erecta* (Der), *D. ananassae* (Dan), *D. willistoni* (Dwi), *D. mojavensis* (Dmo), *D. virilis* (Dvi) and *D. grimshawi* (Dgr). RNA secondary structures between the docking site and selector sequence for exon 4.11 were previously identified[1]. RNA secondary structures between the docking site and selector sequence for exon 4.8 were predicted only in melanogaster group species and housefly. The sequences that make up the core of the stem are highlighted in green.

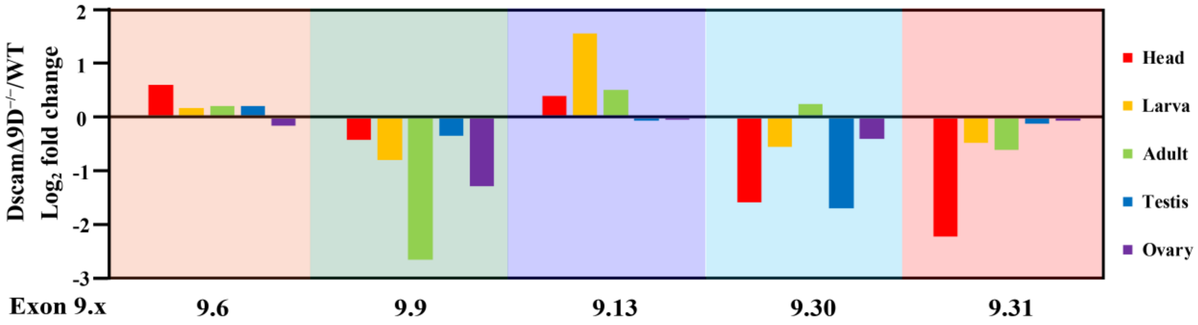


**Figure S3. Deletion of the docking site led to altered frequency of the exon 9 variants**. The $\log_2$ fold change in the frequency of most dominantly expressed variable exon 9s (exon 9.6, 9.9, 9.13, 9.30, 9.31) in various developmental stages and tissues. A relatively similar change trend of exon 9 usage was observed in various tissues of DscamΔ9D$^{-/-}$, although the alterations were subject to developmental and tissue-specific regulation.

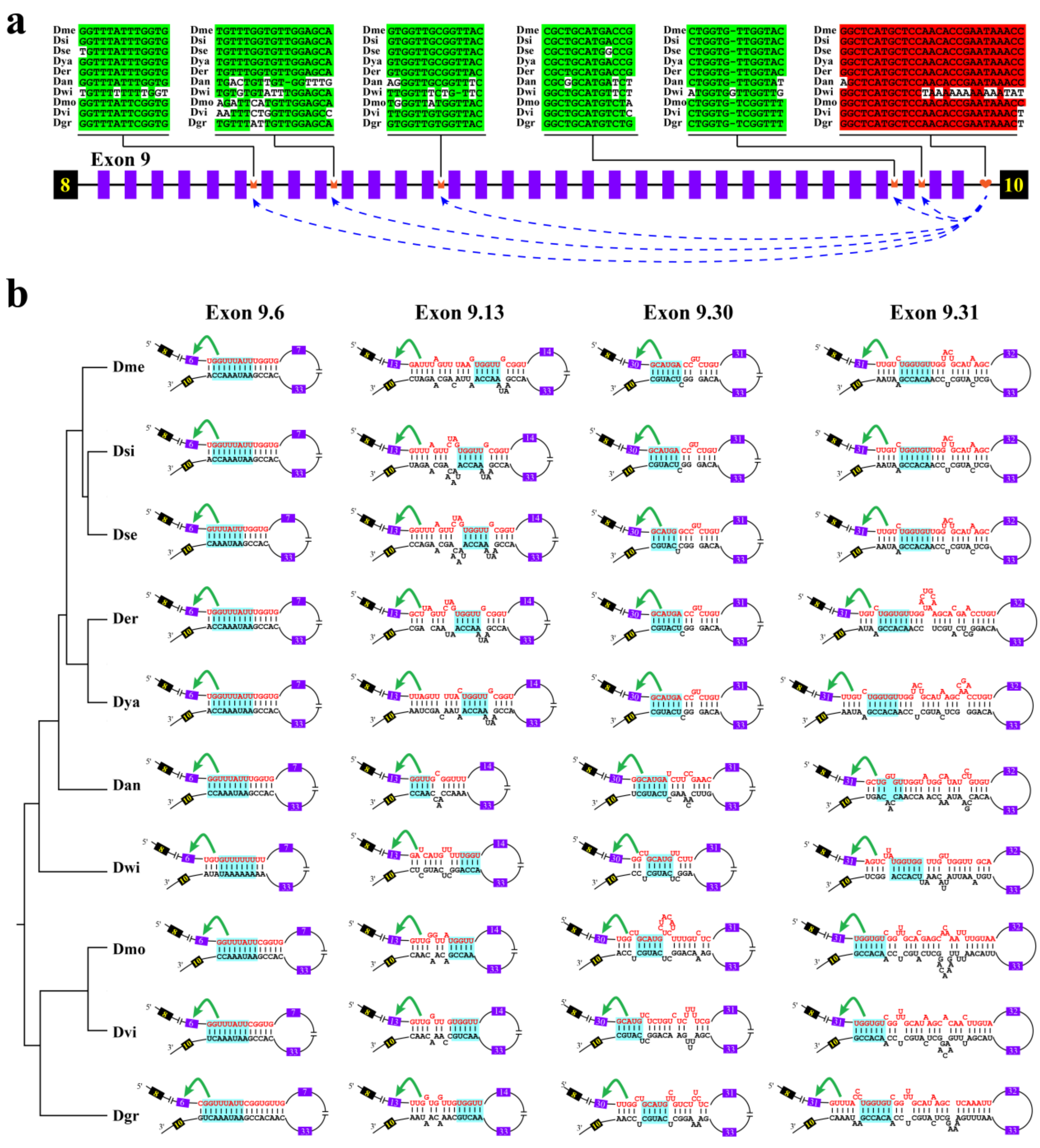


**Figure S4. RNA secondary structures between the docking site and selector sequence are conserved.** (**a**) Overview of the arrangement of the docking site and selector sequence of exon 9 cluster of *Drosophila Dscam1*. Symbols used are the same as those in Fig. 1. Constitutive exons (in black boxes), alternative exon 9s (in purple boxes), docking sites (marked by hearts) and selector sequences (marked by crowns) are shown. Above are sequences of consensus intronic elements for different species. Those *Drosophila* species include *D. melanogaster* (Dme), *D. simulans* (Dsi), *D. sechellia* (Dse), *D. yakuba* (Dya), *D. erecta* (Der), *D. ananassae* (Dan), *D. willistoni* (Dwi), *D. mojavensis* (Dmo), *D. virilis* (Dvi)

and *D. grimshawi* (Dgr). The most identical nucleotides with respect to the docking sites and selector sequences are shaded in red and green, respectively. **(b)** The base pairings between the docking site and selector sequence are conserved in exon 9.6, 9.9, 9.13, 9.30, 9.31 across *Drosophila* species. RNA secondary structures between the docking site and selector sequence for exon 9.9 were previously identified[1]. The base pairings between the docking site and the selector sequences were predicted using the Mfold program[2]. The sequences that make up the core of the stem in *Drosophila* species are highlighted in light blue.

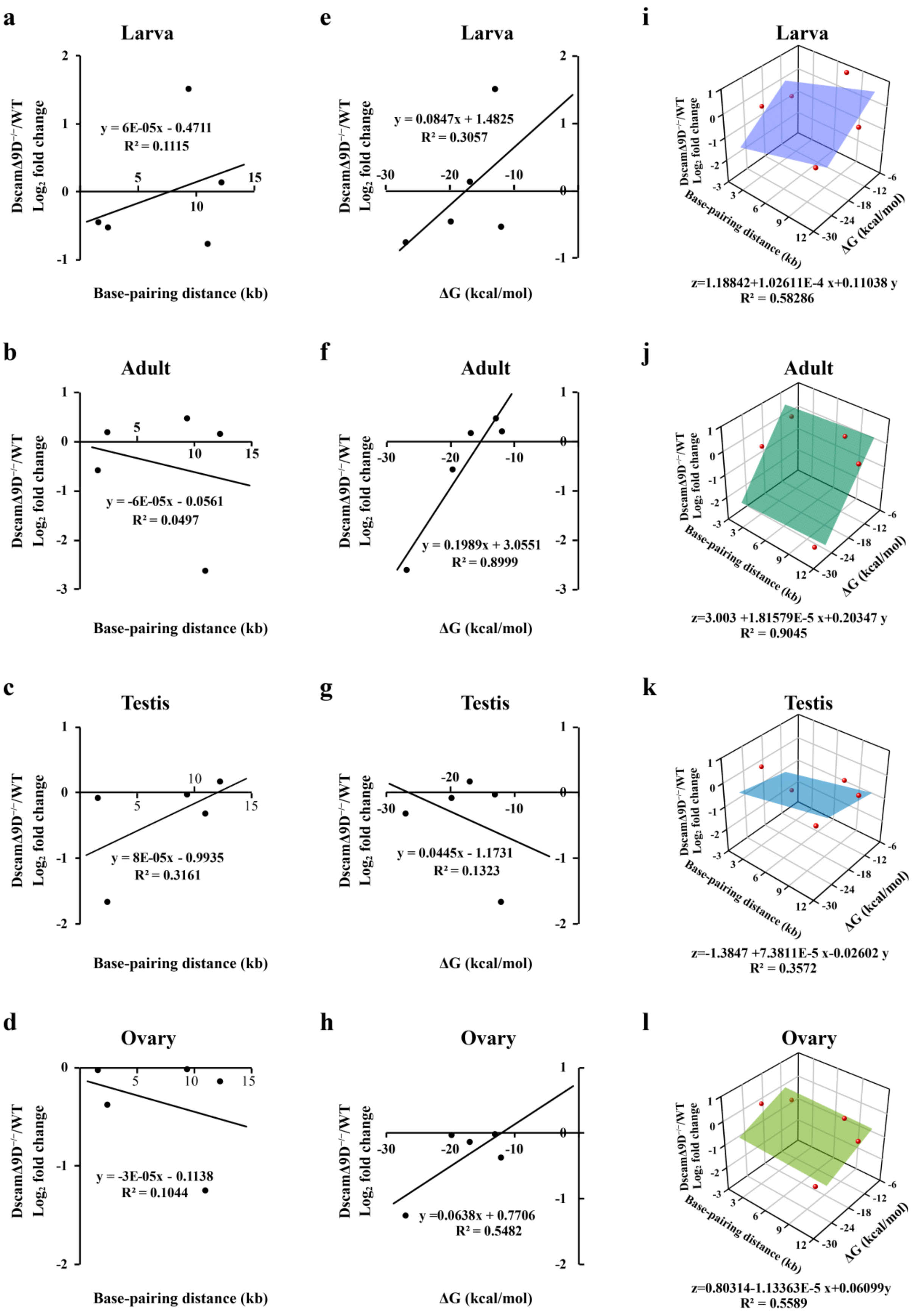


**Figure S5. Deleting the docking site affected the frequency of exon 9 inclusions through disturbing the base pairing interaction in proximity- and strength-mediated manner.**

**(a–d)** Correlation analysis between the distances of the docking site-selector interaction and the fold change in the frequency of the exon 9 variants. The distance of the docking site-selector interaction is plotted as a function of the fold change in the frequency of inclusion of the exon 9 variants in larva (a), adult (b), testis (c), ovary (d). **(e–h)** Correlation analysis between the base pairing strength（ΔG kcal/mol） of the docking site-selector interaction and the fold change in the frequency of the exon 9 variants. The strength of the base pairing between the docking site and selector sequence is plotted as a function of the fold change in the frequency of inclusion of the exon 9 variants in larva (e), adult (f), testis (g), ovary (h). **(i–l)** Multiple linear regression analysis. The combined effect of the distance and strength of the predicted docking site-selector sequence interaction is plotted as a function of the fold change in the frequency of inclusion of the exon 9 variants in larva (i), adult (j), testis (k), ovary (l). These analyses revealed a much stronger correlation of the fold change in exon 9 inclusion with the combined effect of both the distance and strength of the docking site-selector base pairing than with either its strength or distance.

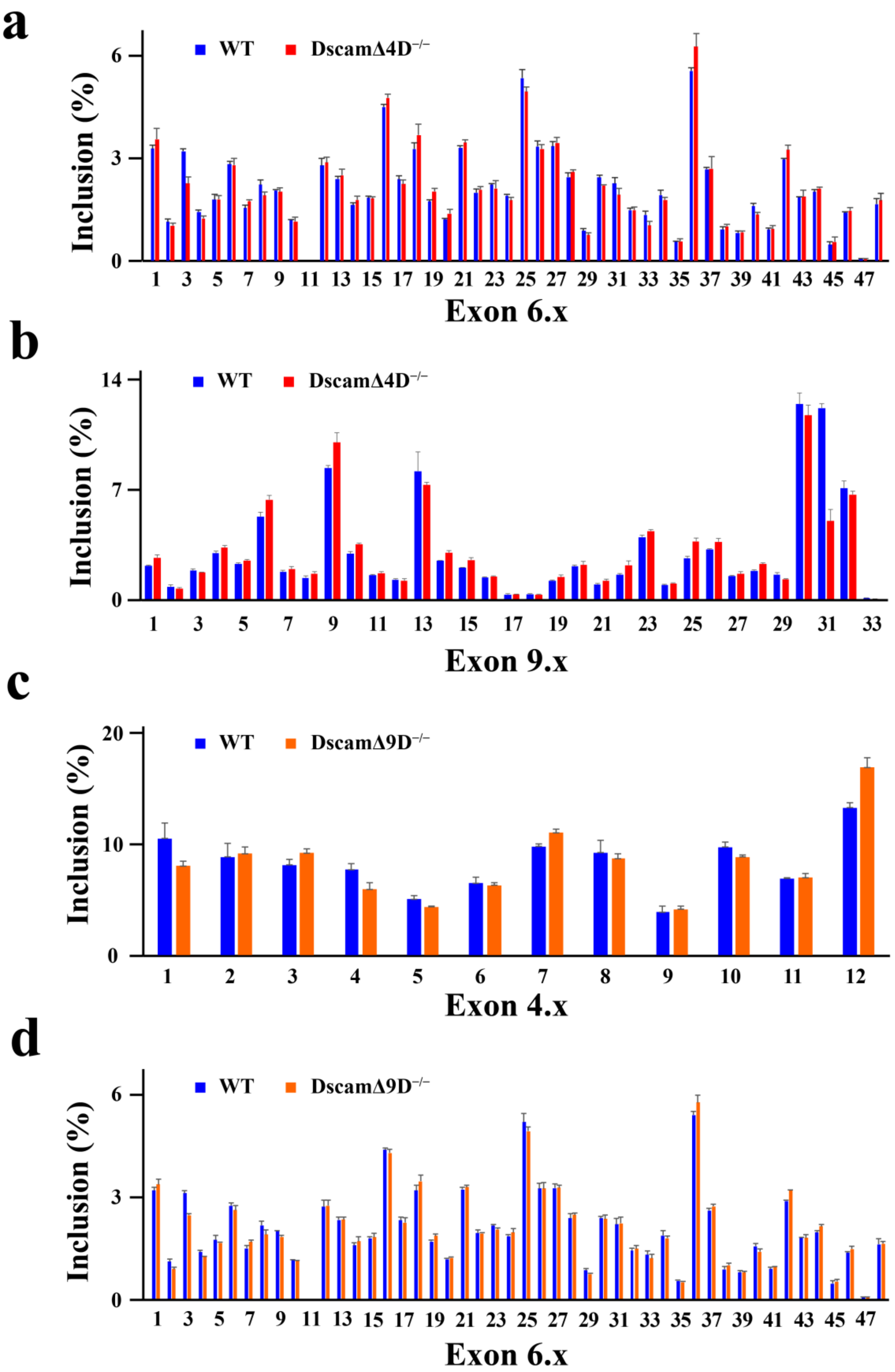


**Figure S6. DscamΔ4D$^{-/-}$ or DscamΔ9D$^{-/-}$ flies show largely independent splicing between exon clusters 4, 6, and 9. (a)** No significant differences in alternative exon 6s inclusion between control and DscamΔ4D$^{-/-}$ flies. **(b)** No significant differences in frequency of most of alternative exons 9 between control and DscamΔ4D$^{-/-}$ flies were observed. **(c)** No significant differences in alternative exon 4s inclusion between control and DscamΔ9D$^{-/-}$ flies. **(d)** No significant differences in alternative exon 6s inclusion between control and DscamΔ9D$^{-/-}$ flies. These results indicate that alternative splicing between different clusters of *Dscam1* is largely independent.

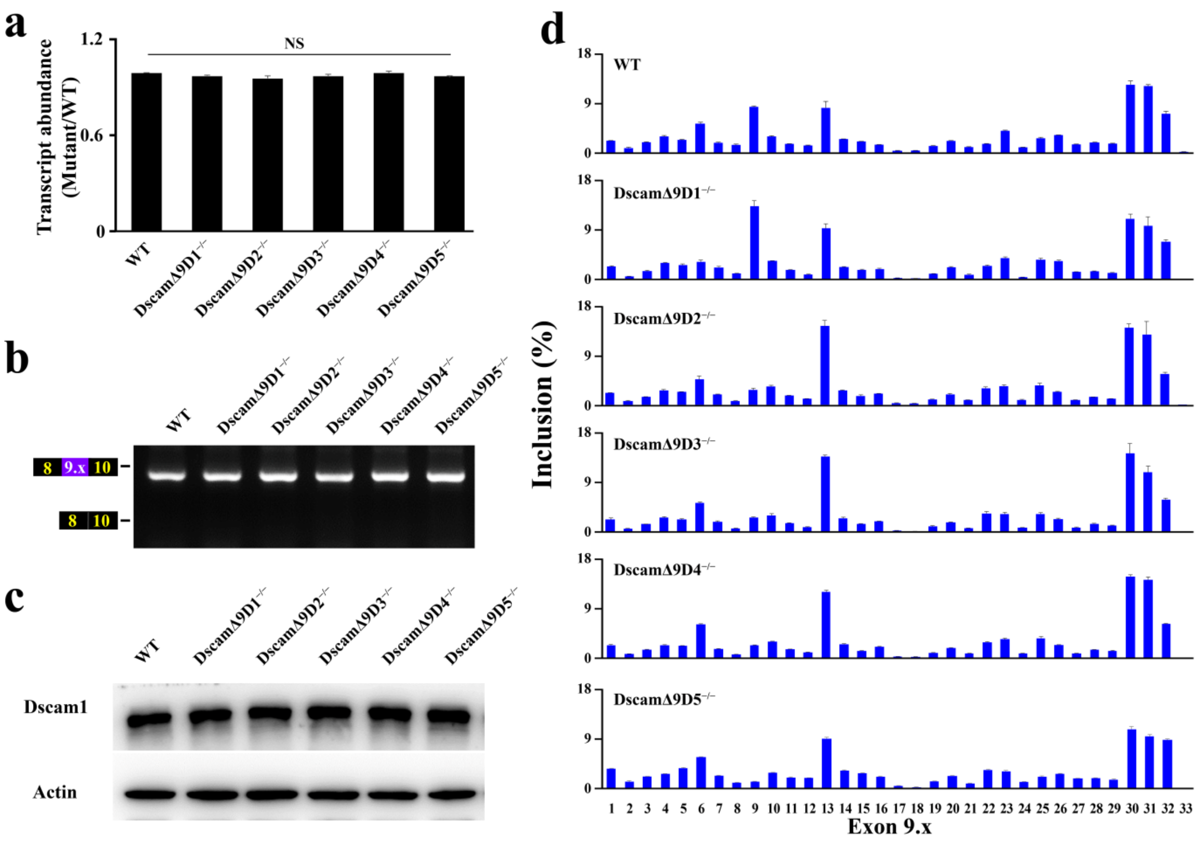


**Figure S7. Molecular characterization of DscamΔ9D1–5$^{-/-}$ alleles. (a)** The transcript level for each *Dscam1* mutant was indistinguishable from wild-type control; Data are expressed as mean ± s.d. from three independent experiments. *P<0.05; **P<0.01; ***P<0.001; NS, not significant (Student's t-test, two-tail). (**b)** The exon 9 inclusions for each *Dscam1* mutant were indistinguishable from the wild-type control. (**c)** Disruption of the docking sites did not affect Dscam1 protein levels. (**d)** Deletion of the docking site dramatically altered the choice frequency of the exon 9 variants. The relative frequency of the exon 9 inclusion in wild type and mutants was shown (from exons 9.1 to 9.33).

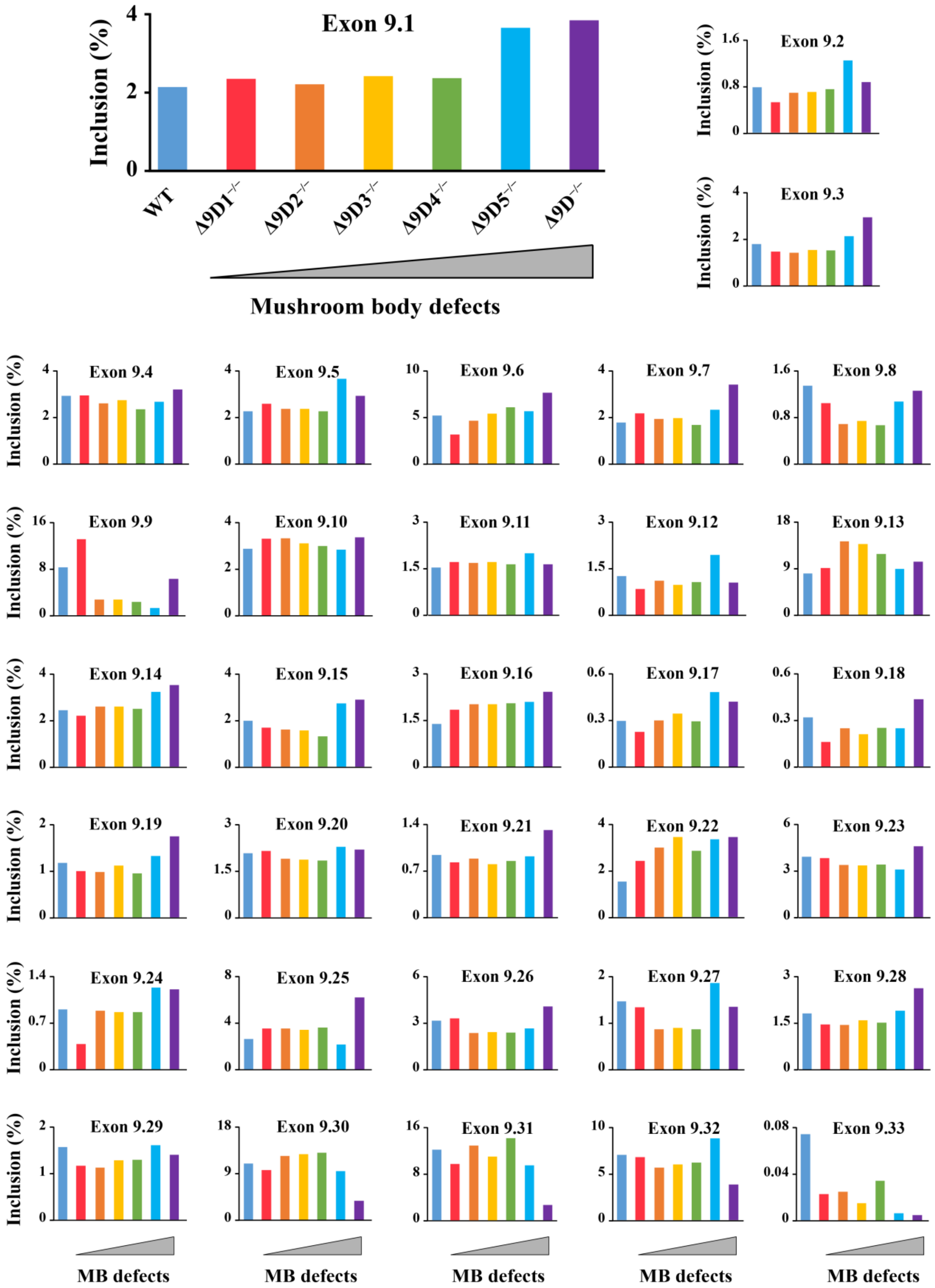


**Figure S8. Analyses of correlation of the expression of specific exon 9 variants with mushroom body (MB) defects.** The histograms show the relationship between inclusion of exon 9 and the proportion of mutant mushroom body defects. Different colors represent different mutants (light blue: wild type, red: DscamΔ9D1$^{-/-}$, orange: DscamΔ9D2$^{-/-}$, yellow: DscamΔ9D3$^{-/-}$, green: DscamΔ9D4$^{-/-}$, sky blue: DscamΔ9D5$^{-/-}$, purple: DscamΔ9D$^{-/-}$). Mutants were arranged from low to high proportion of mushroom body defects. In order to show the inclusion of each exon 9 variants in different mutants, exons 9.1 to 9.33 are represented by different graphs. As shown in the figure, the inclusions of almost all exons have been changed.

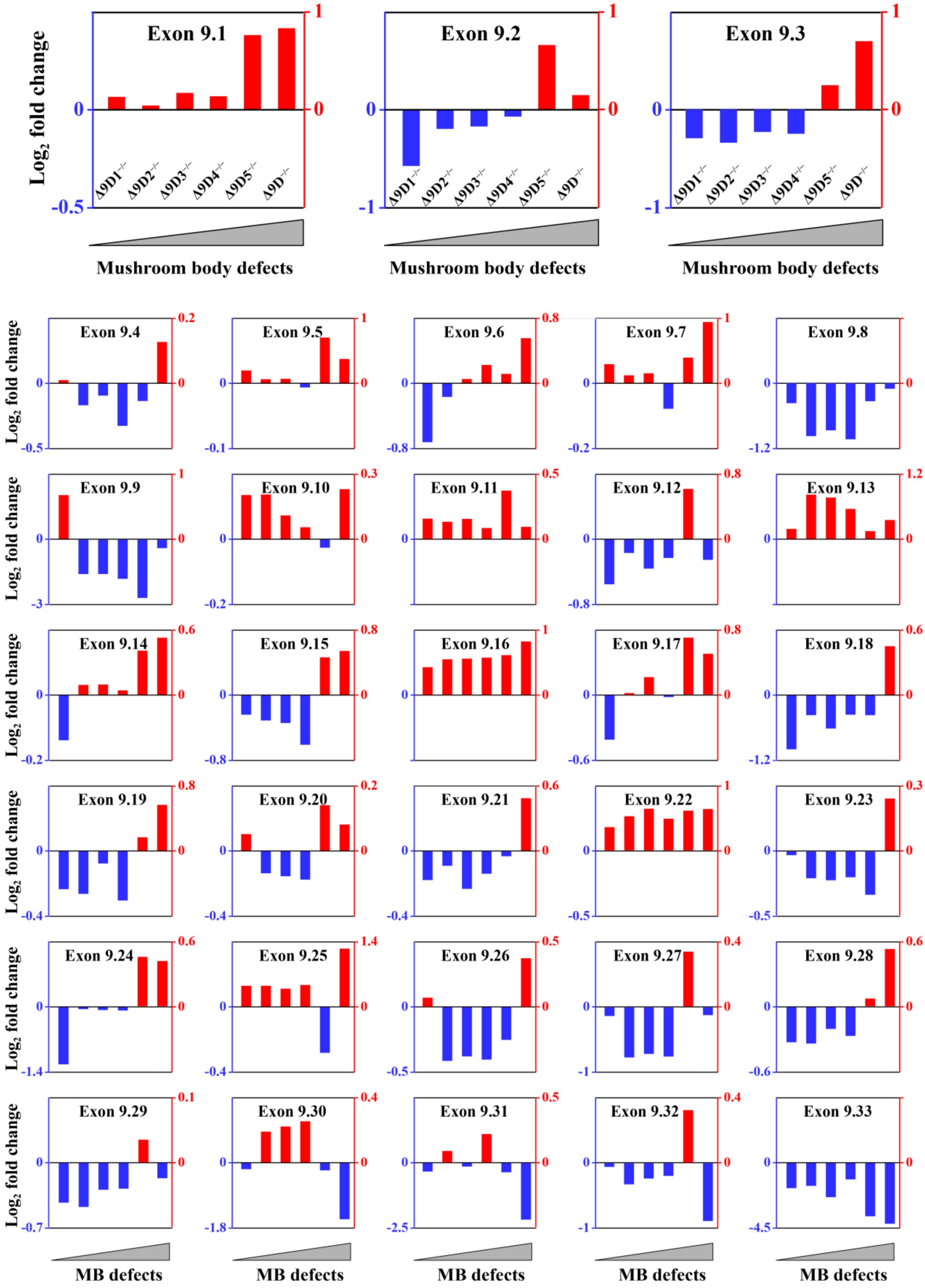


**Figure S9. Analyses of correlation of the expression changes of specific exon 9 variants with mushroom body (MB) defects.** The $\log_2$ fold change of each mutant construct compared to wild type is represented as column chart. Mutants were arranged from low to high proportion of mushroom body defects. In order to show the changes in each exon 9 variants with different mutants, exons 9.1 to 9.33 are represented by different graphs. As we can see from the figures, the inclusions of almost all exons have been changed.

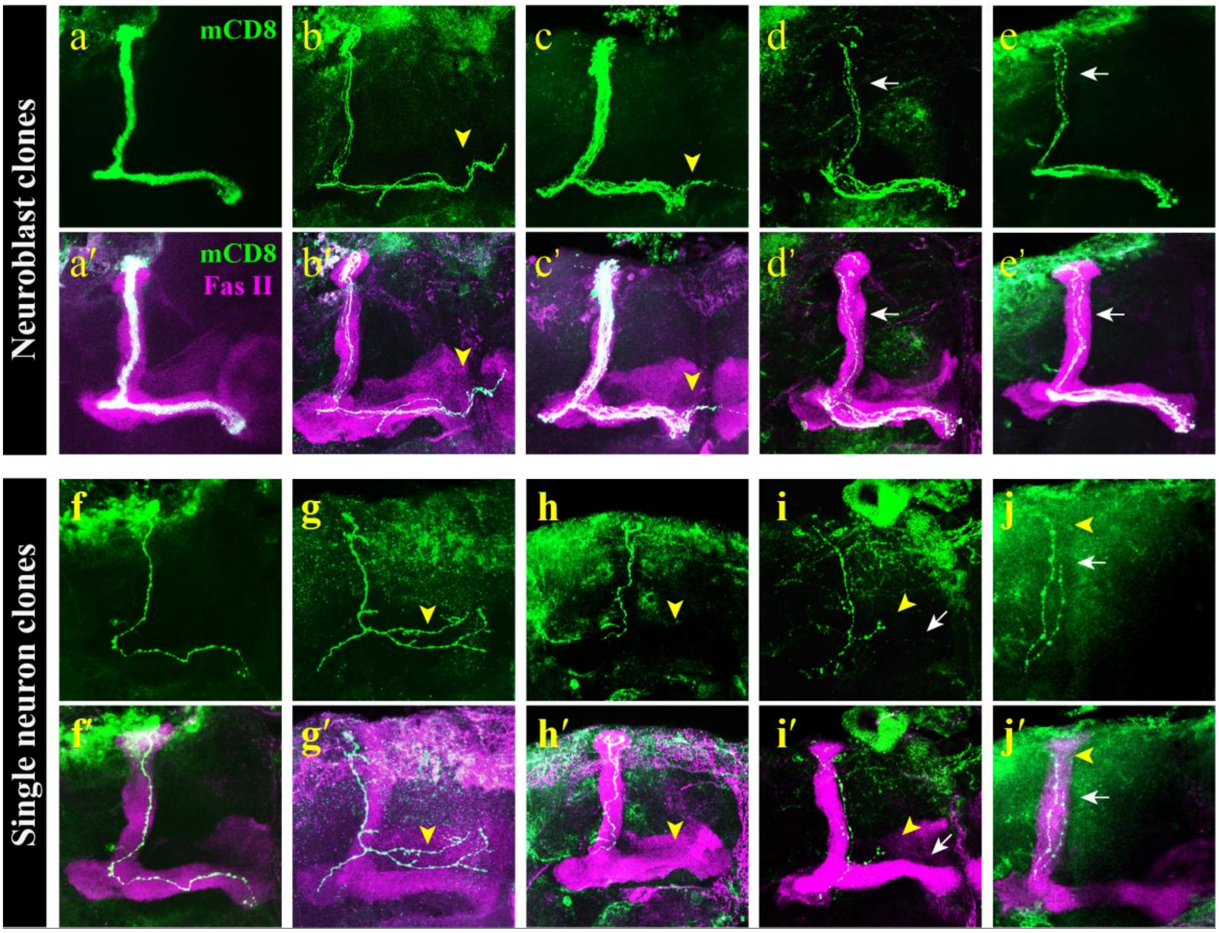


**Figure S10. Phenotypic analysis of *Dscam1* mutant mushroom body axons at neuroblast and single-cell clones (supplemental to Figure 7c–f', h–k').** All drawings and images (a–j') are adult mushroom body of brain hemisphere. a-j are the same as a'-j', respectively, showing FasII staining. The control neuroblast clones show the bifurcated axon with a dorsally and a medially running branch, while DscamΔ4D$^{-/-}$ and DscamΔ9D$^{-/-}$ mutant clones exhibited either a growth defect or a guidance axonal defect, or sometimes a combination of the growth and guidance defects. **(a–e')** Wild type and mutant mushroom body axons in neuroblast (NB) clones (supplemental to Figure 7c-f'). Compared with wild-type neuroblast (NB) clones (a, a'), mutant NB clones display overextension of axon branches (b, b', c, c'), thinning and thickening of α/β axon branches (d, d', e, e'). Yellow arrowhead indicated growth defect that axon branches extended beyond the ends of β lobes (b, b', c, c'). White arrowhead indicated guidance defect that axons in α lobe was thinner than in wild-type control (d, d', e, e'). **(f–j')** Wild-type and mutant mushroom body axons in single neuron clones (supplemental to Figure 7h-k'). Unlike wild type single neuron clones (f, f'), mutant clones exhibited either growth (g, g', h, h',), or a combination of growth and guidance defects (i, i', j, j'). Supernumerary branches (g, g') or an unusual single-branch phenotype with no axon bifurcation at the peduncle end (h, h') were shown in some single-cell clones (yellow arrowhead). In panel i and j, yellow arrowhead indicates growth defect, while white arrowhead represents guidance defect.

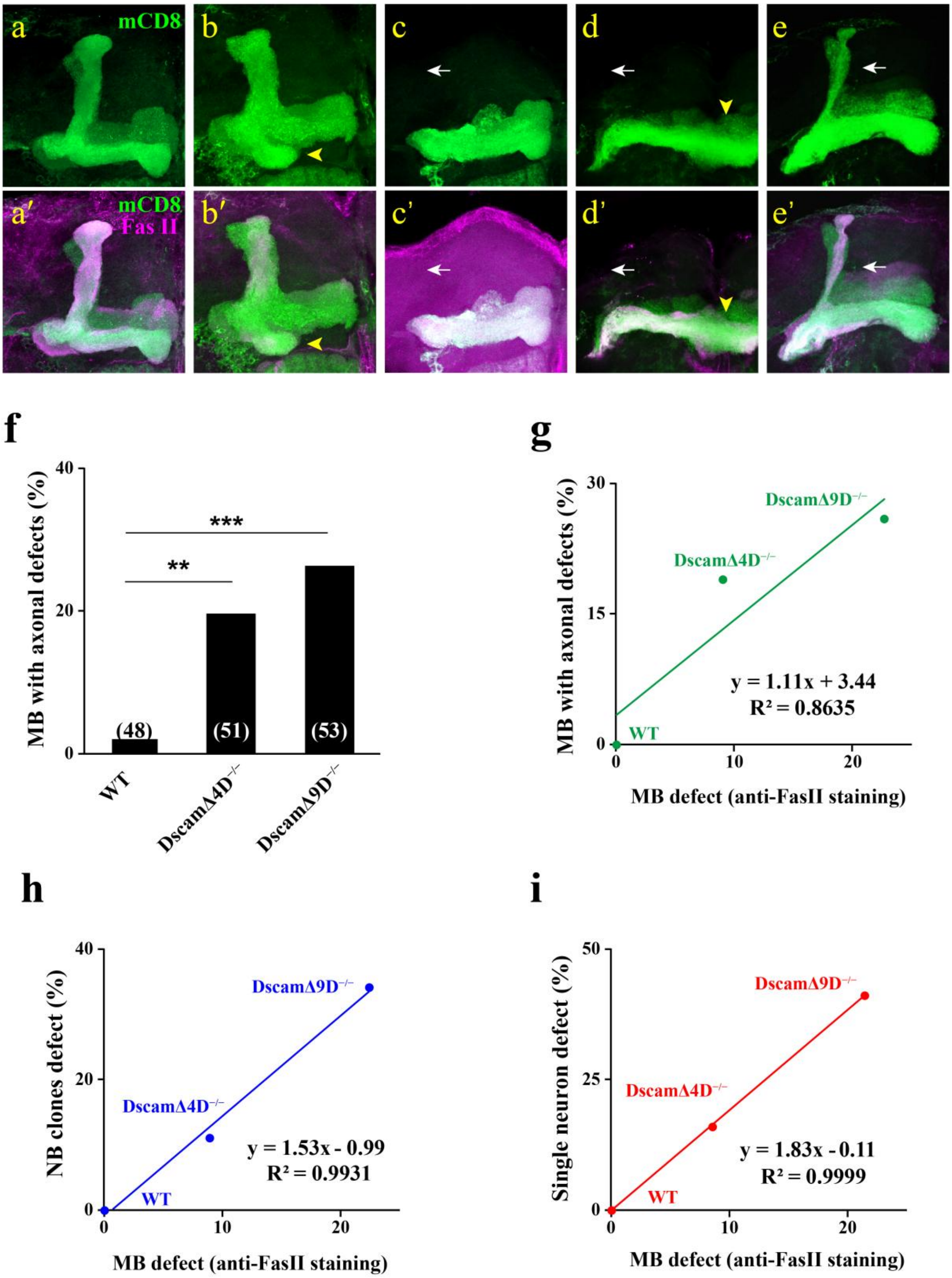


**Figure S11. Phenotypic analysis of *Dscam1* mushroom body axons in isogenic mutant brains. (a–e')** Wild type and mutant axons in large neuroblast (NB) clones. All drawings and images (a–e') are adult mushroom body of brain hemisphere. a-e are the same as a'-e', respectively, showing FasII staining. A control neuroblast clones show the bifurcated axon with a dorsally and a medially running branch, while DscamΔ4D$^{-/-}$ and DscamΔ9D$^{-/-}$ mutant clones exhibited either a growth defect or a guidance axonal defect, or sometimes a combination of the growth and guidance defects. Compared with wild-type neuroblast (NB) clones (a, a'), mutant NB clones display growth (b, b'), and guidance (c, c', e, e') defects, or sometimes a combination of them (d, d'). Yellow arrowhead indicated growth defect that axons did not reach the ends of α/β lobes in some mutant clones (yellow arrowhead, b, b', d, d'). White arrowhead indicated guidance defect that axons in α lobe was thinner than in wild-type control or lost (white arrowhead, c, c', e, e'). **(f)** Quantification of mushroom body axonal defects in isogenic mutant brains. **(g–i)** Correlation of mushroom body defects detected by anti-FasII staining with axonal defects observed in isogenic mutant brains (g), neuroblast clones (h), or single neuron clones (i).
